# Neural replay is connected to latent cause inference and supports fast generalization

**DOI:** 10.64898/2025.12.12.693963

**Authors:** Fabian M. Renz, Shany Grossman, Nathaniel D. Daw, Peter Dayan, Christian F. Doeller, Nicolas W. Schuck

## Abstract

Dynamic environments require generalizing knowledge through two computational mechanisms: Inferring clusters of equivalent observations and propagating reward expectations within a cluster. While neural replay has been linked to both, past work has not studied these processes simultaneously, leaving their relative importance for replay unclear. We used fMRI and computational modeling to investigate latent cause inference and replay in 52 participants during a value-based task with shared rewards across bandits. Participants learned this structure and achieved one-shot generalization after reversals, consistent with a latent cause inference process. fMRI during predecision pauses revealed backwards replay of shared-reward bandits in the visual cortex and medial temporal lobe. Trial-wise fluctuations in visual replay strength and content were better explained by value updates than by structure learning. Conversely, abstract reward structure representations were localized specifically to the MTL. These results demonstrate that on-task replay serves reward propagation within learned latent structures to facilitate fast generalization.

## 1 Introduction

A hallmark of adaptive behavior is the ability to generalize from past experiences to guide decisions in novel situations. This capacity relies on representations that capture the underlying decision-relevant structure of the environment while filtering out details irrelevant for reward prediction (Shepard, 1987; Niv, 2019; Kaplan et al., 2017). When environments are novel or changing, successful generalization therefore requires to first infer an environment’s hidden task states, a process often formalized as latent cause inference (LCI), which leverages similarities across observed events to infer the shared, unobservable causes that generated them (Collins and Frank, 2013; Lloyd and Leslie, 2013; Gershman et al., 2015). Having established that distinct observations arise from the same latent cause, one can begin to generalize information (e.g. value) from one observation to the other.

A central question is how the brain initially learns and subsequently leverages such latent structure for flexible generalization. Recent work suggests a critical role for neural replay, the sequential reactivation of neural states corresponding to past experiences (Foster, 2017; Wittkuhn et al., 2021; Schuck and Niv, 2019). However, the specific computational function of replay in learning and leveraging hidden structure remains debated, with research pointing to two distinct roles. One line of work suggests replay serves to update value expectations or plan immediate actions. For instance, rodent studies have shown that replay sequences often depict future paths to remembered goals, predicting immediate behavioral choices (Pfeiffer and Foster, 2013). In this view, replay utilizes the inferred structure to propagate reward information to related states, performing “non-local” value updating (Mattar and Daw, 2018; Liu et al., 2021b). In contrast, alternative accounts of replay argue that replay is primarily concerned with memory, and in particular building and maintaining the abstract representation of the environment itself (Ólafsdóttir et al., 2018). This view is supported by findings that replay often represents past goals or non-local areas rather than the immediate future path, decoupling it from current choices (Gupta et al., 2010; Gillespie et al., 2021), as well as by evidence for structure replay in tasks that do not involve reward learning (Wittkuhn et al., 2025). Notably, Gillespie et al. (2021) found that replay often “lags” behavioral learning, suggesting it may function to consolidate the structure of the task, stitching together separate experiences to form a coherent map rather than drive immediate planning Ou et al. (2025); Bakermans et al. (2025). Recently, these views have recently been framed as complementary processes, where replay might prioritize either map-building or planning depending on the uncertainty of future goals (Sagiv et al., 2025; Ólafsdóttir et al., 2018).

Although methodological advances progressively allow the detection of replay in humans (e.g., Wittkuhn and Schuck, 2021; Wittkuhn et al., 2025; Liu et al., 2021b, 2019), empirical evidence distinguishing between these computational roles remains scarce, particularly during the initial discovery of task structure. Recent magnetoencephalography (MEG) work has shown that replay of unobserved paths can support value generalization across shared reward structures (Liu et al., 2021b). This finding aligns with the “non-local” value updating account (Mattar and Daw, 2018), suggesting that replay utilizes offline periods to propagate new reward information across learned structure. Yet, because prior work has largely relied on pre-learned associations, it remains unknown how replay interacts with the discovery or adaptation of that structure. A critical mechanistic question persists: Does replay actively drive the inference of latent causes or does it primarily exploit an emerging structure to propagate value?

To capture the dynamical interaction between structure learning and value generalization and to identify its potential neural bases, we combine behavioral, neuroimaging, and latent cause inference modeling to investigate how latent structure is learned and used in the service of generalization. On the first day of our experiment, participants learned a novel graph structure linking multiple visual sequences to shared or independent reward outcomes, which repeatedly reversed throughout time (similar to Liu et al. 2021b). On the second day, we changed the latent reward structure, requiring flexible reorganization for appropriate generalization. This design allowed us to track both the acquisition of structure and its adaptation to change. To capture the intertwined structure and reward learning processes, we fitted a latent cause inference model to participants’ data and, following our previous work (Schuck and Niv, 2019; Wittkuhn and Schuck, 2021; Wittkuhn et al., 2025), measured replay using functional magnetic resonance imaging (fMRI). Results indicate that participants learned the structure of our task and achieved successful generalization in the second half of day 1. A latent cause model fitted human behavior better than standard reinforcement learning models, successfully capturing structure acquisition and generalization processes. Neural analyses in visual cortex revealed that replay was tightly aligned with participants’ knowledge of the latent structure: its strength increased as the reward relationships were learned on day 1 and adapted after the structural change, including a pronounced shift in directionality for the newly generalizable path. These dynamics indicate that replay is closely coupled with the inferred structure and supports value updating, consistent with our LCI model in which trial-wise non-local value updating of the unobserved path was the strongest predictor of replay. Moreover, we found that multivariate patterns in the medial temporal lobe reflected the emerging reward structure and reorganized following structural change.

## 2 Results

### Structure learning facilitates fast value generalization

Fifty-two healthy participants (mean age 27.4; 25 females) completed a 4-session fMRI experiment across two consecutive days (Fig. 1D). The primary purpose of our study was to investigate the role of neural replay in (a) learning the latent task structure and (b) the structure-enabled value generalization.

**Figure 1:**
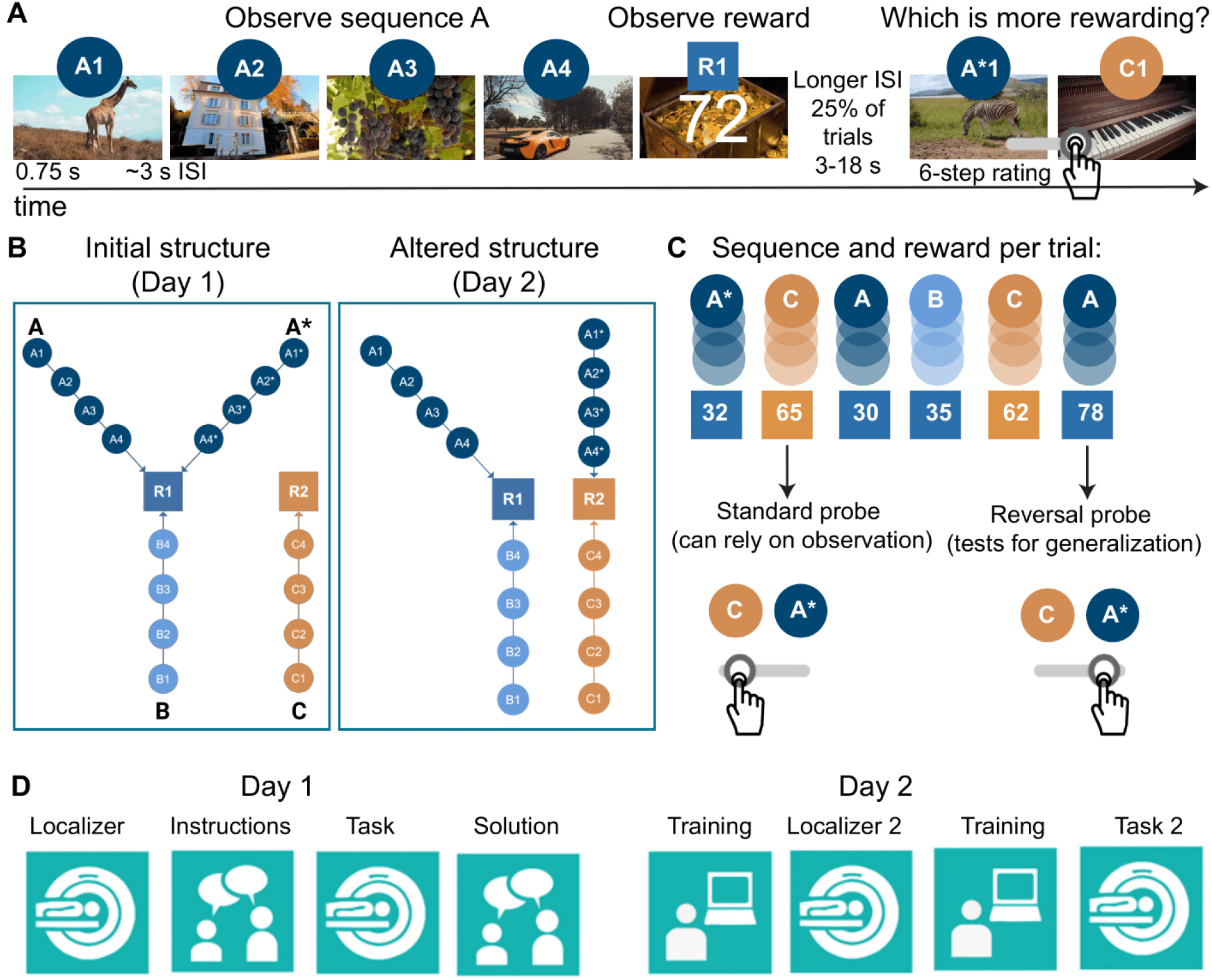
Task design and experimental paradigm. (A) Each trial presented a sequence of four 750 ms video stimuli leading to a reward outcome (points), and was then followed by a choice between two options using a 6-point confidence slider. (B) On Day 1, three sequences (A, A*, B) led to a shared reward outcome (R1) and sequence C to an independent outcome (R2); A and A* shared the semantic categories for each sequence position (e.g. animals: giraffe and zebra). On Day 2, after 40% of the trials, sequence A* covertly switched from R1 to R2, requiring participants to learn the new underlying structure and adapt their generalization strategy accordingly. (C) Trials were either *standard trials*, in which values could be judged from direct observation, or *reversal trials*, which tested generalization to unobserved sequences after a reward reversal. Correct structure knowledge enabled generalizing value changes from an observed sequence (e.g., A) to others sharing its reward contingency (e.g., A* and B). (D) Experimental timeline: Each day began with an fMRI localizer used for classifier training, followed by task training and the main task. Day 1 involved structure discovery from experience, ending with explicit revelation of the structure. Day 2 therefore began with full structure knowledge, allowing assessment of exploitation and adaptation to the mid-session structure change.

In both days of the main task, participants had to decide between four sequential bandits (A, A^∗^, B and C), each consisting of a sequence of four 750 ms videos that led to a reward outcome (e.g., A1 → A2 → A3 → A4 → R1; videos depicted distinct objects e.g., giraffe, grapes, basketball; see Fig. 1A and SI Fig. 4). Each sequence terminated in a scalar outcome (0–100 points), with rewards for three bandits (A, A^∗^, and B) sampled from one latent reward distribution (R1) and rewards for bandit C sampled from a second distribution (R2); these distributions were noisily sampled on each trial and periodically reversed such that which latent reward was currently higher changed over time. The video sequences of bandits A and A* depicted objects from the same categories (e.g., A1 and A^∗^1 depicted a giraffe and a zebra, i.e. both animals; the other matched categories were houses, cars, and fruits), whereas the content of all other videos was unrelated. On each trial, participants first passively observed one of the four video sequences and its outcome, and were then asked to choose between two video screenshots using a preference slider (1-6). The choice options were always matched in their positional distance from the reward (e.g. B2 vs. C2, see Methods for details). Feedback on overall accumulated reward was provided every 10 trials. The task involved 120 trials structured into 5 blocks, and reward outcomes reversed episodically every 15-20 trials (i.e., 6 reversals in total, see Methods).

As expected, participants successfully learned reward contingencies and adjusted their choices upon reversals to the bandits with the highest outcomes (see Fig. 2A; remaining reward schedules in SI Figure 1), demonstrated by an increase in accuracy from 68.7 % accuracy in Block 1 to 84.8 % in Block 5 (linear effect of block in a linear mixed effect model: *β* = 0.041, *z* = 6.46, *p* < .001, see Fig. 2B).

**Figure 2:**
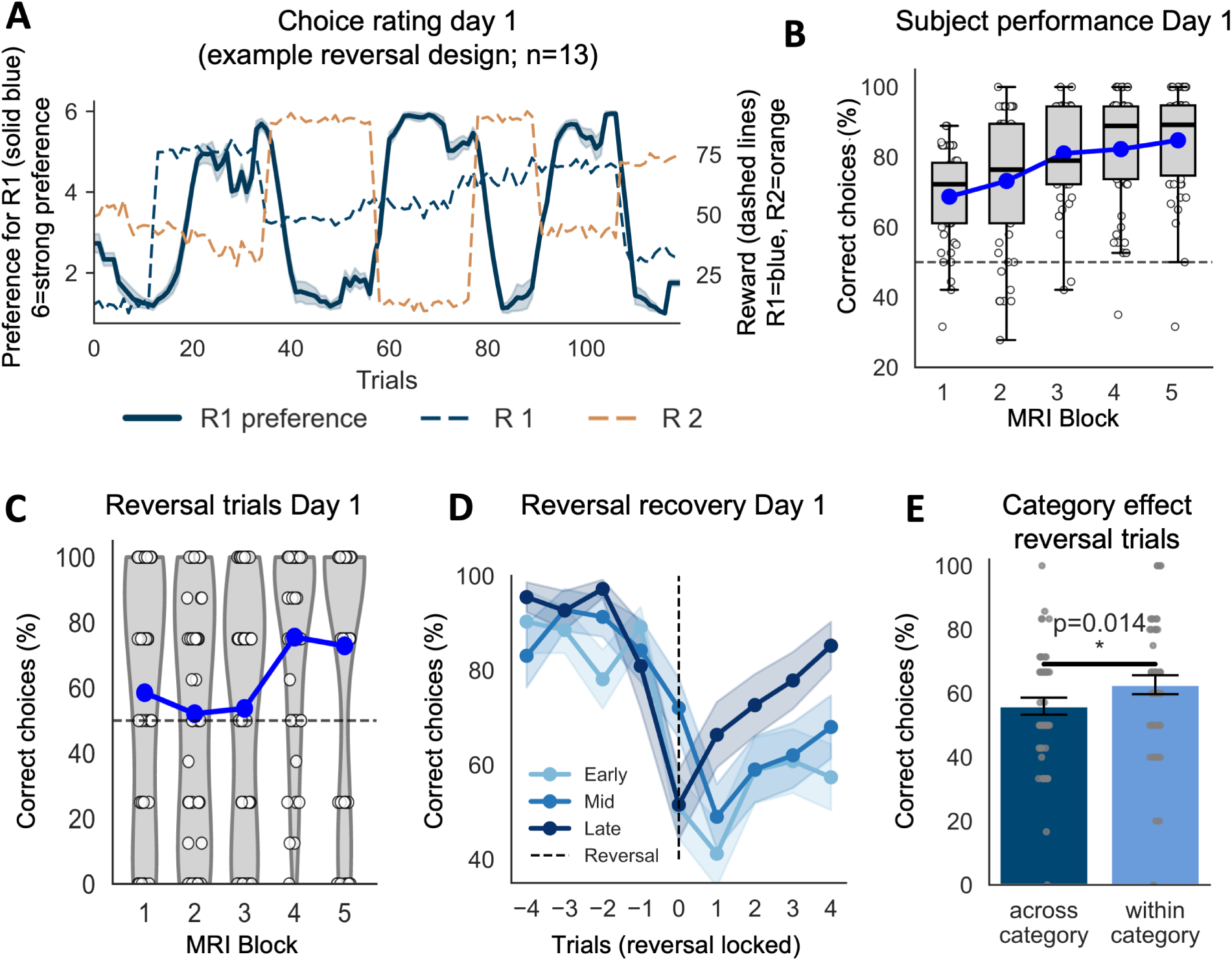
Behavioral performance on Day 1 demonstrates structure learning and adaptation. (A) Example preference ratings for one reward schedule (solid blue line; 1–6 scale) for R1 compared with the true reward values (dashed lines; R1=blue, R2=orange) across Day 1. Participants successfully adapted to the reward reversals: when R1 > R2, ratings for R1 increased, indicating a strong preference, whereas when R2 > R1, ratings for R1 decreased, reflecting a shift in preference toward R2. (B) Accuracy across all trials, showing progressive improvement over the course of Day 1. (C) Performance on reversal trials that required generalization to unobserved sequences. Accuracy begins at chance early in training and increases with experience, reaching above-chance levels in blocks 4 and 5. (D) Recovery from reversals, time-locked to the reversal onset (trial 0), shown for early (reversals 1 and 2), mid (reversals 3,4), and late reversals (reversals 5,6) across Day 1. Recovery becomes faster over the course of training, indicating improved use of latent structure. (E) Semantic similarity effect on Day 1 reversal trials: performance was higher when generalizing within category-sharing sequences (e.g., observing A and being probed on A*).

A core feature of our task on Day 1 was that, unbeknownst to participants, the outcomes of three sequential bandits (A, A^∗^ and B) were sampled from the same latent mean (shared reward, R1), while bandit C was associated with an independent reward (R2, unique reward, Fig. 1B). Our main behavioral question was whether participants would learn about the correlation structure of the rewards and use it to generalize outcome observations among all three related bandits. To test this, we used trials immediately after a reward reversal to provide a primary measure for participants’ knowledge of the graph structure. Specifically, we asked participants to choose between two nodes of the graph that had not been observed since the reversal. To answer correctly in these reversal-locked trials, participants had to generalize value from an observed bandit (e.g. A) to one of the correlated bandits (e.g., A^∗^), based on shared reward contingencies rather than direct observation (henceforth reversal trials, see the right choice in Fig. 1C). Note that in these trials, responding based on one’s memory of a bandit’s last observed reward would lead to objective accuracy of less than 50%: Although the last rewards observed for bandits A* and C indicated that C is better than A*, for instance, in reversal trials participants should prefer A* over C because they had just observed that A resulted in a higher outcome on the last trial.

An analysis of reversal trials on Day 1 showed that participants improved accuracy from around 55% in blocks 1-2 to 74% in blocks 4-5 (Fig. 2C, main effect of block in lme: *β* = 0.052, *z* = 3.041, *p* = 0.002; performance in blocks 4 and 5 was above chance, both p *<* 0.05, FDR corrected). This pattern was also evident in the trial-wise accuracy following reward reversals (Fig. 2D). During early reversals (first 2), accuracy showed a maximum drawdown to 41% immediately after the reversal and did not recover to reliably above-chance performance within the subsequent generalization trials (post-reversal trial 4: 60.8%, *p* = 0.02; trial 5: 58.0%, *p* = 0.11). In the middle phase of the task (reversals 3 and 4), the maximum drawdown decreased to 49%, following which participants returned to above-chance performance by the fifth reversal generalization trial (post-reversal trial 4: 62%, *p* = 0.106; trial 5: 68%, *p* = 0.009). During the late stage of the task (reversals 5 and 6), the maximum drawdown was 51%, after which participants performed above chance from the second post-reversal trial onward (post-reversal trial 2: 67%, trial 5: 84%; all *p <* 0.001 from trial 2 onward, FDR-corrected; Fig. 2D). The slight dip before the reversal likely reflects participants beginning to adapt as rewards start changing but have not yet reversed, as reversals transition through 1–2 intermediate steps between stable reward levels (see Methods). As mentioned above, videos associated with bandits A and A^∗^ were semantically similar, while videos in A/A^∗^ and B were not. We hypothesized that semantic similarity would facilitate generalization (e.g., Tompary and Thompson-Schill, 2021). Indeed, reversal trials requiring generalization between category-sharing sequences (e.g., from A to A^∗^) showed superior performance compared to cross-category generalization (e.g., from B to A^∗^; paired t-test *t*(50) = −2.56, *p* = 0.014, see Fig. 2E). Hence, while generalization occurred for bandit pairs with and without semantic similarity, such similarity did enhance structure learning and generalization.

We also included 10 trials that probed two options that led to the same reward (e.g., A2 vs. A*2, both leading to R1), which therefore had no “correct” solution under the initial structure. On these trials, participants showed significantly weaker preferences than on trials involving options from different reward sources, which elicited more extreme ratings. (A linear mixed-effects model predicting rating extremity (center = low, extreme = high; trial type *β* = 0.343, *z* = 13.26, *p* < .001).

### A latent cause inference model captures structure discovery and generalization

One mechanism that can explain structure learning and value generalization is a process known as latent cause inference (LCI). This framework posits that the brain infers hidden, unobservable causes to explain the structure of observed patterns of events and then generalizes on the basis of inferred latent causes (Gershman et al., 2010; Lloyd and Leslie, 2013; Gershman et al., 2015). To test this idea, we developed a LCI model of our task that used a Chinese Restaurant Process prior with particle filtering to perform approximate inference (see Sanborn et al., 2010, see Methods) about hidden causes that might have generated the observed video sequences and their rewards. Different particles represent alternative hypotheses about latent cause structures and are resampled according to how well they account for the observed data. Because the model continuously inferred the structure of latent causes given past observations, it could learn about reward correlations and generalize values after reward changes by discovering shared latent causes across paths (Fig. 3A). We applied the LCI model (4 free parameters: concentration parameter *α*, categorical update *β*, hazard rate *h*, and choice temperature *θ*) to data from our task and tested it against three alternative models: a simple temporal difference (TD) model that learned about each bandit independently (two free parameters learning rate *α*, temperature *θ*), and two versions of ‘clairvoyant’ TD models that applied each TD update to all bandits which shared outcomes with either a single or differential learning rates (the models are clairvoyant because they magically know from the start how outcomes can be generalized). The TD model had the worst model fit (AICc = 69.6), due to its failure to generalize following reversal trials (see SI). The clairvoyant TD models had marginally better model fits because they did generalize (single learning rate AICc = 63; differential learning rates AICc = 56,6), but their model fit was limited by their inability to capture the learning progression throughout the first day (as well as the structure change during the second day, see below). The LCI model’s flexibility in both discovering and updating structure yielded the best fit (AICc = 43.8; Fig. 3D). Inspecting the LCI model more closely confirmed that it captured key behavioral phenomena across both days (e.g., reversal trial generalization, see Fig. 3B; reversal recovery SI Fig. 2C).

**Figure 3:**
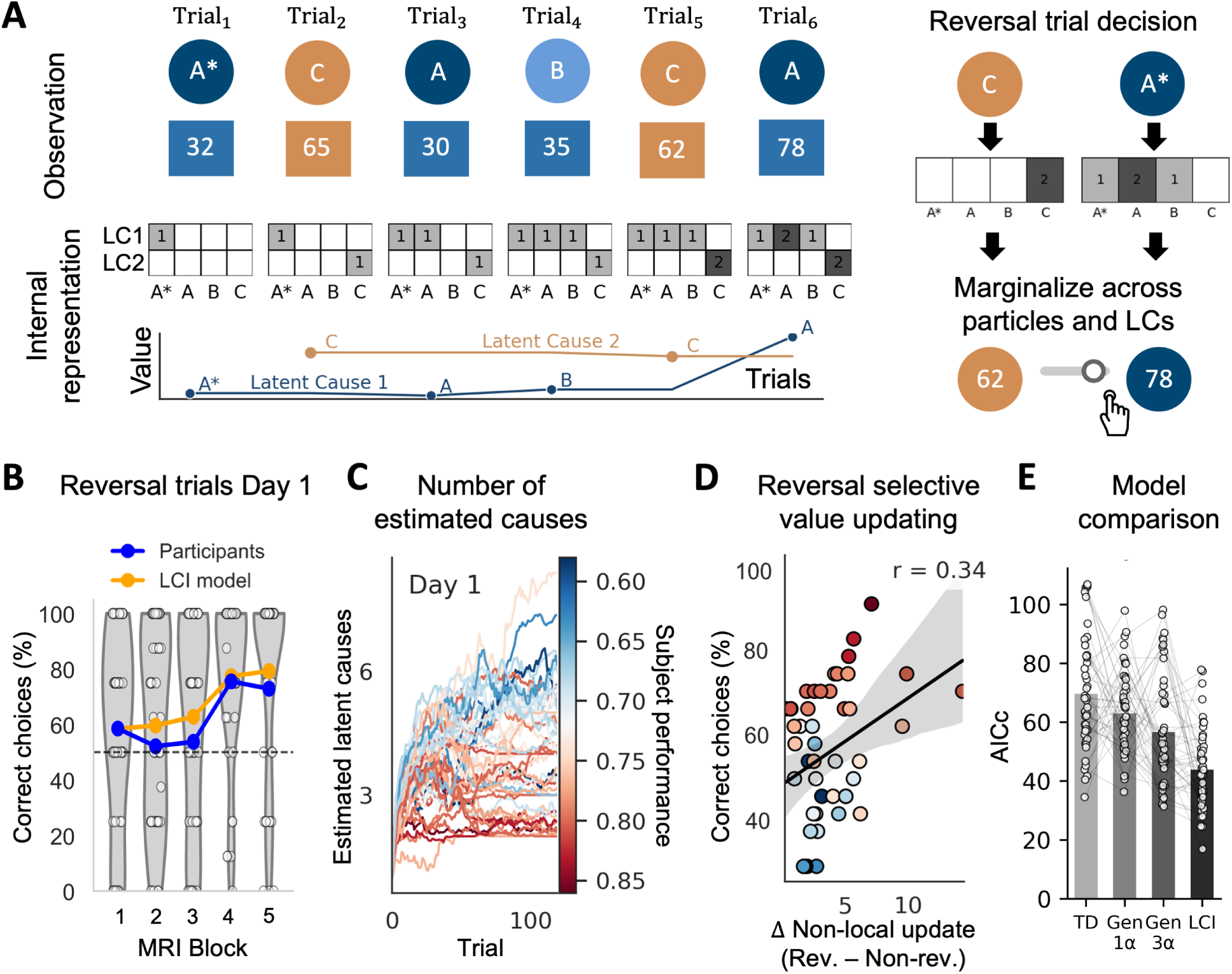
Latent cause inference model explains structure learning on Day 1. (A) Illustration of the latent cause inference model. On each trial, the model computes the likelihood of the current observation for all existing versus a potential new latent cause, combining (i) how often the observed sequence has previously been assigned to that cause and (ii) the cause’s current reward expectation. The likelihood is combined with a Chinese Restaurant Process prior, which favors reuse of frequently inferred causes. Particles then branch over all possible assignments and are resampled in proportion to how well each branch accounts for the observed data. As sequences become associated with shared latent causes, updated values assigned to one sequence can be accessed by other sequences through their shared latent-cause representation, enabling generalization. Trial-by-trial value estimates are obtained by marginalizing over latent causes and particles. (B) Reversal trial performance calculated as percent correct choices on Day 1. The LCI model captures the gradual improvement in participants’ generalization accuracy across blocks. (C) Individual differences on Day 1. Participants with higher behavioral performance were best fit by models inferring fewer latent causes, consistent with more efficient structure discovery and generalization through shared latent causes. (D) Individual differences in selective value updating relate to reversal performance. The x-axis shows the model-derived increase in non-local value updating for unobserved sequences during reversal trials relative to non-reversal trials (Rev. − Non-rev.). The y-axis shows participants’ reversal performance, calculated as the percentage of correct responses. Greater model-derived selective value updating was associated with higher behavioral performance. Point color reflects overall task accuracy, with warmer colors indicating higher performance (same color scale as panel C). (E) Model comparison across the full experiment. The LCI model outperforms both temporal-difference models lacking generalization and models with fixed, non-adaptive generalization structure. This performance difference is preserved when the analysis is conducted separately for each day (see SI Fig. 2A).

The latent cause model also captured individual differences in structure learning. Participants with higher overall accuracy were fit by models discovering fewer latent causes (Fig. 3C; linear mixed-effects model: *β* = −0.022, SE = 0.006, *z* = −4.03, *p <* 0.001), indicating efficient structure discovery (i.e., value generalization was achieved by grouping sequences with shared rewards under a common latent cause). Moreover, the selectivity of value updating within the model correlated with participants’ reversal performance: participants’ accuracy, calculated as the percentage of correct choices in reversal trials, was positively associated with the degree to which the model exhibited higher non-local updating during reversal compared to stable periods. (LME: *β* = 0.021, *z* = 2.825, *p* = 0.005; Fig. 3D). Together, these results suggest that statistical learning supports the inference of latent structure that scaffolds value generalization, enabling selective, non-local value updating when contingencies change while preserving stability during steady states.

### Backward replay selectively reactivates unobserved generalizable paths

Following previous work (Liu et al., 2021b), our core neural hypothesis was that value generalization was facilitated by neural replay, i.e. that generalization from one bandit to another was driven by post-outcome backward replay of the bandit to which outcome knowledge was generalized. In line with our behavioral findings, we specifically expected such “non-local” replay of unobserved sequences to occur for bandits that shared reward contingencies. To test this hypothesis, we inserted 18-second breaks after reward presentation, just before the decision phase, i.e. the moment when value generalization would be needed to support subsequent choices. These prolonged inter-stimulus intervals (ISIs) were distributed throughout the task but had systematically higher frequency after trials following reward reversals, as these were the critical moments during which non-local updating was most needed for successful decision-making (40 per session). Almost all reversal trials were associated with prolonged ISIs and we did not observe any performance difference comparing non-reversal trials with and without prolonged ISI (*t*(50) = 1.393, *p* = .170).

To decode replay-related neural signals, we first trained logistic regression classifiers on fMRI patterns from independent localizer sessions conducted before and after the task (sessions 1 and 3, see Methods and SI Fig. 3). Classifiers were trained in a one-versus-rest fashion to distinguish between the 16 individual video stimuli. We first focused on a visual ROI (object-selective regions in the ventral stream, as in Wittkuhn and Schuck (2021), for details see Methods), which yielded a low between-class confusion (see Fig. 4A) and very high classification performance (mean probability of true class at peak: 71.2%, *t*(49) = 44.02, *p <* .001, Fig. 4B) that is a prerequisite for subsequent replay analyses. Note, one classifier was excluded from all replay analyses because it showed elevated activation during the reward stimulus (see Methods), which could have biased the prolonged ISI. Importantly, the exclusion of this classifier did not selectively impact any condition. Because condition assignments depended on the presented sequence, the sequence containing the excluded classifier was represented in all four conditions introduced below (see Fig. 4D).

**Figure 4:**
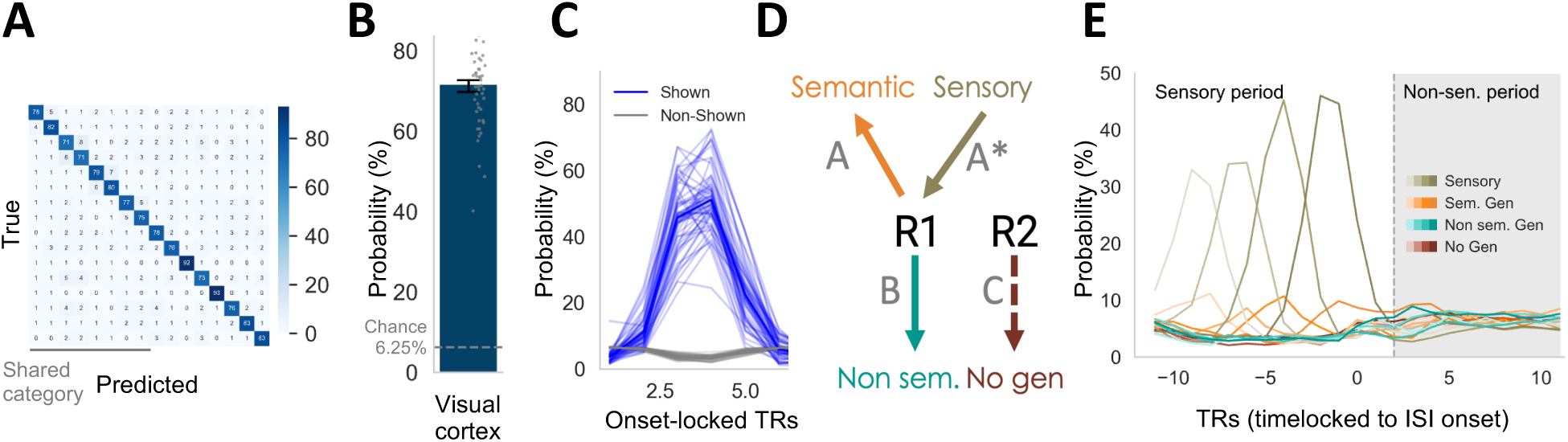
Training and applying classifiers. (A) Confusion matrix from localizer sessions showing accurate decoding of 16 video stimuli in visual cortex with minimal category bias. (B) Cross-validated localizer performance in visual cortex (avg. probability of true class 71.2%; chance level 6.25%, dashed line). (C) Transfer to the main task: classifier probabilities peak following stimulus onset for shown items (blue) but not non-shown items (gray). (D) Replay conditions. After viewing sequence A* (sensory, olive), unobserved sequences were classified as semantic generalization (A, shared reward and category, orange), non-semantic generalization (B, shared reward only, turquoise), or no generalization (C, unrelated, brown). (E) Averaged classifier probability time courses aligned to prolonged ISI onset (time 0). Strong sequential sensory responses precede ISI onset, followed by sequential reactivation during the non-sensory period. The sensory period refers to the initial TRs dominated by stimulus-driven responses, whereas the non-sensory period begins once these sensory responses return to baseline.

Classifiers trained on localizer sessions also successfully decoded the displayed stimuli during the main task (probability at peak in the visual ROI: 51.1%, *t*(50) = 31.35, *p <* .001, Fig. 4C and SI Fig. 5). To study replay during the ISI periods, we employed the sequential slope ordering analysis (SODA) method developed and used in Schuck and Niv (2019); Wittkuhn and Schuck (2021); Wittkuhn et al. (2025). The main idea of SODA is that sequential neural activity will lead to a specific pattern of ordered classifier probabilities across time, which can be detected by analyzing the slopes of TR-wise regression coefficients that test a hypothesized sequential order. Hence, we could test for instance whether class probabilities of classifiers of the A branch were ordered as expected (*p*(A1) *> p*(A2) *> p*(A3) *> p*(A4)), and similarly for the other three sequences. The slope assesses average ordering, and thus does not guarantee that every comparison is conjunctively significant. We excluded the reward state from the sequence modeling, as it was shared across all four sequences. Given the set of possible within-branch orderings, we constructed for each ISI period four sequence slope models and categorized them according to the branch that had been observed in the trial immediately before that ISI (Fig. 4D). First, sensory model coefficients referred to the branch just seen before the ISI and hence reflected sequential activation potentially arising due to the actually observed video sequence. Second, the semantic generalization model quantified sequential reactivation of the bandit that is semantically related to, and shares a latent reward with, the bandit shown before the ISI (e.g., reactivating the sequence A4→A3→A2→A1 after observing A^∗^), if any; Third, non-semantic generalization model slopes reflected reactivation of sequences that share the same latent reward, but not the semantic category (e.g., replaying the sequence B4→B3→… after observing A^∗^). Finally, the non-generalization model captured reactivation of sequences that are unrelated to the just observed one (e.g., C4→C3→… after observing A^∗^). We hypothesized that we would observe Semantic and Non-Semantic generalization effects that indicate backward replay. Sensory forward activation elicited by the presented video sequence served as a sanity check for our analyses, and Non-generalization sequences functioned as a control condition.

Focusing on the visual ROI, we found strong evidence for sensory-driven sequential activation during the presentation of the sequence and the early TRs of the prolonged ISI, as expected (Fig. 4E). As shown in Fig. 5A, the regression slope testing the sensory sequence showed a sinusoidal pattern, with positive coefficients (indicating forward ordering, *β*s *>* 0 at TRs -11–6 relative to the ISI onset, all *p*s *<* 0.01, corrected) followed by negative coefficients (reflecting the expected sign flip that is part of any forward sequence, see (Wittkuhn and Schuck, 2021), *β*s *<* 0 at TRs -4–1, all *p*s *<* 0.01, corrected). Hence, our classifiers were able to detect objective sequential activation.

**Figure 5:**
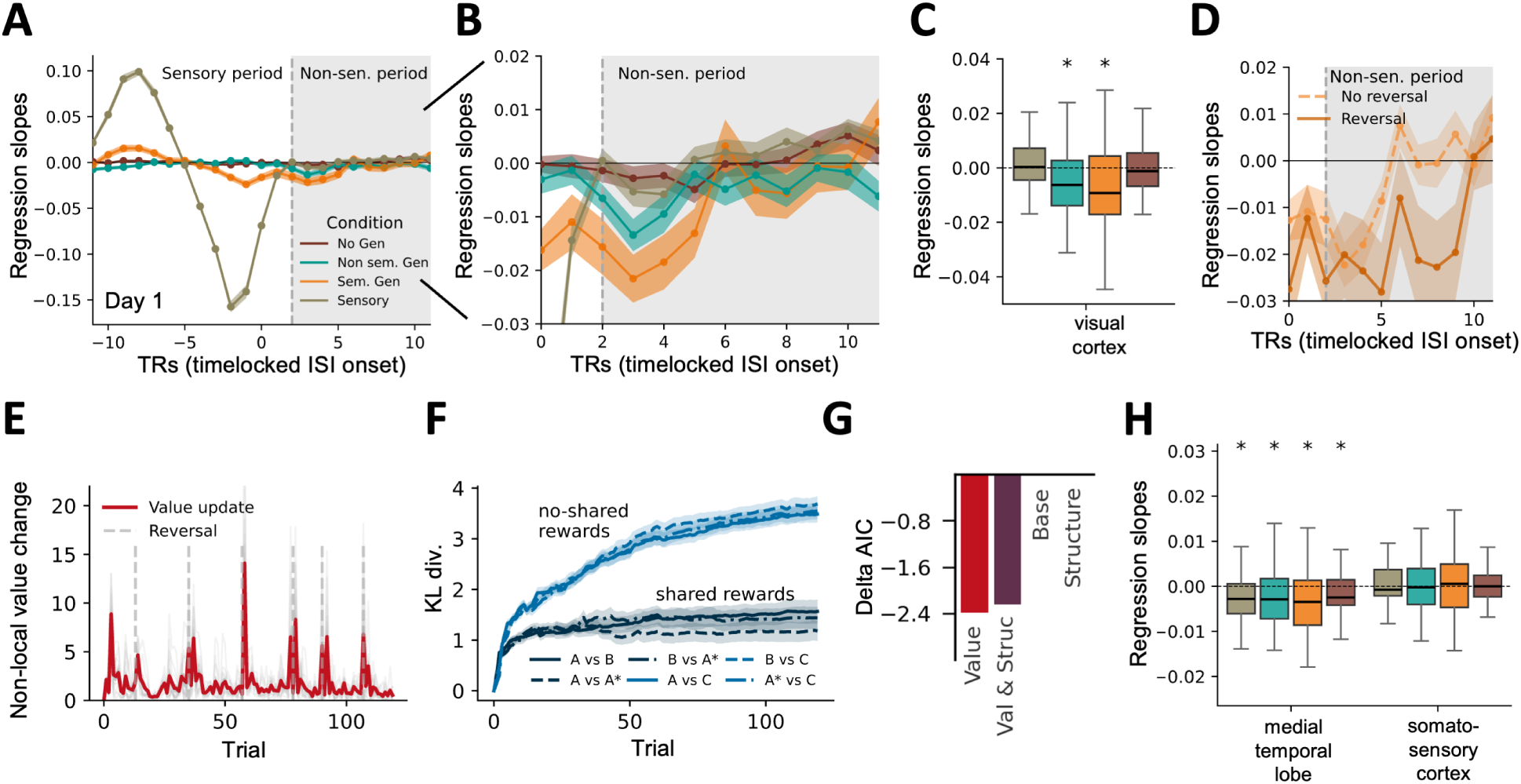
Detecting and relating replay to value updating and structure learning. (A) Regression slope time-courses. Sensory sequences show forward then backward patterns during stimulus presentation. After the sensory effect subsides (gray shaded area), reward-linked sequences (semantic, non-semantic) exhibit selective backward replay, while unrelated sequences do not. (B) Zoomed regression slope timecourses during the non-sensory period, high-lighting distinct reactivation for semantic and non-semantic generalization. (C) Aggregated regression slopes during the non-sensory period. Semantic (orange) and non-semantic (turquoise) sequences show selective backward replay. Error bars show SEM. (D) Semantic generalization slopes split by reversal vs. non-reversal trials. Stronger reactivation occurs following reversals, consistent with need for non-local value updating. (E) Latent cause model–derived nonlocal value updating for unobserved options peaks around reward reversal trials. The red line shows the mean trialwise change in unobserved sequence values, averaged across participants with the same reward schedule. The predictor used in the linear mixed-effects model was computed at the subject, trial, and sequence level, capturing the model-derived value update for each sequence (for example, the update to B after observing A on that trial) and relating the magnitude of this update to the amount of reactivation evidence observed for the corresponding sequence. (F) Latent cause model structure learning, quantified as the evolving similarity between paths over time (symmetric KL divergence between their latent-cause probability distributions). Paths that shared rewards showed increasing similarity (decreasing KL divergence), whereas non-shared paths remained distinct. The corresponding predictor in the linear mixed-effects model was the trialwise change in this KL divergence metric, capturing moment-to-moment updates in inferred task structure. (G) Trial-by-trial updating of value predicts replay reactivation beyond a baseline model including only condition as a predictor. (H) Aggregated regression slopes during the non-sensory period for MTL and somatosensory cortex. In the MTL, all conditions show significant backward replay across sequences, albeit weaker than in visual cortex. In contrast, the somatosensory cortex (control) shows no evidence of systematic replay for any condition.

We also observed that unobserved but semantically related categories co-activated during the stimulus presentation period, although confusion among these semantically related categories during the localizer was minimal (Fig. 5A). To test whether this effect indeed reflected meaningful co-activation or merely classifier uncertainty, we quantified how the classifier distributed residual probability across non-presented classes during the localizer versus main task. For each trial, we computed the difference between (i) the probability assigned to the unobserved but semantically related class and (ii) the mean probability assigned to all other unobserved classes that were not part of shown or related category sequences. If co-activation resulted solely from classifier uncertainty, the magnitude of this difference should remain the same in the localizer and main task, while associative links between category members as reported previously (e.g., Wimmer and Shohamy, 2012) would lead to a larger difference in the main task. A linear mixed-effects model confirmed a significant increase in category-partner bias during the main task relative to the localizer (*β* = 0.009, *z* = 2.65, *p* = .008), indicating that unobserved but semantically related items were activated more strongly than can be accounted for by the baseline classifier confusion.

We next asked whether we could detect replay in the visual ROI during the post-sensory period. We conservatively defined this period as the TRs when the coefficients of the above-described sensory model had returned to 0, which was true starting from 2 TRs into the prolonged ISI (avg. slope: 0.0005; t-test against 0: *t*(50) = 0.233, *p* = .81), see vertical line in Fig. 5A,B). To test whether generalization-related visual replay occurred in this period, we examined the *β* coefficients of *condition* i.e., the semantic and non-semantic generalization models, in comparison to the no generalization and sensory models. A linear mixed effects model of average *β* slopes across the entire post-sensory ISI period with condition as a predictor revealed significant differences across sequence conditions (main effect condition: *z* = 5.89, *p <* 0.001, see Fig. 5C). Post hoc tests showed that both semantic generalization (*t*(49) = −2.55, *p* = 0.028) and non-semantic generalization (*t*(49) = −3.27, *p* = 0.008) exhibited significant negative effects, indicating systematic backward replay. In contrast, no evidence was found for sensory replay (*t*(49) = 0.76, *p* = 0.600) or for no-generalization sequences (*t*(49) = 0.16, *p* = 0.876; Fig. 5C). Direct comparisons further confirmed that sensory replay was significantly weaker than both non-semantic generalization (*t*(50) = 4.10, *p <* 0.01) and semantic generalization (*t*(50) = 3.81, *p <* 0.01). We found no evidence for differences between semantic and non-semantic generalization, either in the overall post-sensory effect (*t*(50) = –0.26, *p* = .80) or at individual TRs. Within the non-sensory window, the smallest uncorrected p-value occurred at TR 5 (*t*(50) = 1.98, *p* = .053). After correcting for multiple comparisons, however, no time point reached significance (all *p >* .24).

Having established that visual replay occurs selectively for generalizable rewards, we next asked whether trial-by-trial fluctuations in replay on Day 1 predicted subsequent choices and generalization. Averaging behavioral accuracy across the following five trials, replay strength predicted better performance (*β* = −0.036, *z* = −2.38, *p* = .017). Replay strength did not predict accuracy on the immediately following decision, however (mixed-effects model: *β* = 0.029, *z* = 1.03, *p* = .302). Given that participants first had to learn about the reward structure before they could begin to generalize, we also tested whether generalization replay increased in strength over time. This was indeed the case (main effect of block *β* = −0.001, *z* = −2.7, *p* = 0.007, across semantic and non-semantic generalization), supporting the idea that the detected visual replay reflects the learned cluster structure. We did not observe any difference between semantic and non-semantic generalization in the increase over time (interaction of block and condition in lme: *β* = 0.002, *z* = 1.59, *p* = 0.112)

Stronger replay for generalizable (unobserved but reward-related) sequences compared to unrelated sequences, increase in selective replay over time, and a link to future performance demonstrate that replay in visual cortex was not driven by recent stimulus exposure, but instead reflected the reward learning based on participants knowledge of the task structure.

### Non-local replay is linked to value reversals

Having established the selective and dynamic nature of visual replay, we sought to determine which factors triggered it. Prior theoretical and empirical work has indicated that a main function of replay could be to facilitate value updating (Liu et al., 2021b; Schaul et al., 2015; Mattar and Daw, 2018; Sinclair and Barense, 2019), and emphasized the effect of prediction errors, which in our study occurred predominantly following reward reversals. Some work has emphasized the role of replay specifically in *non-local* value updating, which uses observed prediction errors to update non-observed states to which values can be generalized (Liu et al., 2021b). By linking three video sequences to the same outcome on Day 1, our task encouraged fast value generalization, such that replay would be expected following reversals from a non-local value updating perspective.

Given that we found no evidence for replay in the non-generalization condition, we focused on modeling the average trial-wise slopes of the semantic and non-semantic replay models (averaged over all post-sensory TRs) as a function of period (5 trials following a reversal, vs. stationary periods, see Methods) and replay condition (semantic vs. non-semantic). Slopes were averaged within each trial across the TRs of the non-sensory period. This analysis showed indeed that reversal periods were a significant predictor of replay (main effect period: *z* = −5.218, *p <* 0.001; Fig. 5D). Model comparison to a baseline model without reversal period also showed significantly improved model fit for the full model with a reversal factor (likelihood-ratio test: *χ*^2^(2) = 19.18, *p* < .001).

### Neural replay aligns with latent cause inference dynamics

While the observation of reversal-triggered replay is interesting, reversals are an experimenter-defined task aspect that participants can only infer indirectly. Hence, the link between task period and replay falls short of providing a link between a genuine cognitive variable and replay. We therefore used the latent cause models that were fitted to participant’s behavior and extracted two cognitive variables of interest that could drive replay, with the goal of providing a statistical model of learning-related trial-by-trial fluctuations in replay. We focused on two main aspects of learning that have been proposed to play a role in replay: nonlocal learning to attribute value to different causes Mattar and Daw (2018), and learning of the latent cause structure itself Sagiv et al. (2025). To test these hypotheses, we used the model to estimate the magnitude of value and structure learning occurring at each trial, and asked whether these correlated with replay. While our model is agnostic to the actual process by which these aspects of learning are implemented in the brain, it served as a way to statistically disambiguate our alternative hypotheses.

We quantified the amount of non-local value change that occurred in each participant’s model following each reward observation, i.e., how much the model updated the value estimate of A after observing B and so forth (Fig. 5E). Contrasting the non-local value updates in our model for reversal trials vs. stationary periods showed that in our model average non-local value updates were larger during reversal periods than during stable periods (*t*(46) = 9.541, *p* < .001), suggesting that this might be a plausible driver of replay. To quantify trial-wise structure learning, we examined how similar the model’s latent-cause probability distributions were across sequences. For every sequence pair and each trial, we computed the Kullback–Leibler (KL) divergence between their latent-cause distributions. Lower divergence indicates that the model assigns similar latent-cause probabilities to both sequences, effectively mapping them onto the same underlying structure. As shown in Fig. 5F, sequences that shared a reward exhibited consistently lower divergence, whereas non-shared sequences showed higher divergence. To align this measure with our trial-resolved value-update analysis, we derived the trial-by-trial change in divergence, capturing how strongly the model updated its structural beliefs on each trial. This trial-wise update served as a proxy for ongoing structure learning, in direct parallel to the model’s non-local value updating.

Having computed these two scores, we asked to what extent they could explain trial-by-trial replay fluctuations in the visual ROI (Fig. 5G). A linear mixed effects model that predicted fluctuations in replay of each non-observed path after each trial showed that including the non-local value change variable as a predictor improved model fit significantly over a baseline model with only a factor for replay condition (AICs: -8375.9 vs. -8373.6; likelihood ratio test value change variable and condition vs. condition only: *χ*^2^(2) = 6.39*, p* = 0.041). Note that the metric we included from the LCI model made predictions not only about fluctuations in overall replay strength, but included separate expectations for each arm, i.e. how strong replay of A* vs. B states was in a given trial, depending on the individual learning history of a given participant up to the trial in question. Using a more generic model that predicted the same replay strength for each arm did not reach statistical significance (likelihood ratio test averaged value change variable and condition vs. condition only: *χ*^2^(2) = 5.488*, p* = 0.064). Including the model-derived measure of structure learning, in contrast, did not improve model fit over the baseline condition-only model either (AIC -8373.6, likelihood ratio test: *χ*^2^(2) = 3.999*, p* = 0.135). Thus, replay in our task seemed mainly related to non-local value fluctuations, which we could predict in a trial and path specific manner from our model. Note that non-local value fluctuations and reversals were correlated, and we did not find evidence that the non-local value model performed better than a model that incorporated only reversal period, i.e. adding non-local value change to a model that already included a regressor for task period did not yield a significant improvement (AICs: -8377.8 vs. -8379.3, likelihood ratio test: *χ*^2^(2) = 2.433*, p* = 0.296).

Lastly, we applied the same decoding and sequentiality analyses to classifier evidence from an anatomically defined medial temporal lobe ROI (MTL; hippocampus, parahippocampal cortex, and entorhinal cortex), motivated by the hippocampal system’s established role in replay in rodents (Wilson and McNaughton, 1994; Skaggs and McNaughton, 1996; Ólafsdóttir et al., 2018) and MEG evidence which appears to indirectly localize human replay signals to hippocampal regions (e.g., Liu et al., 2019, 2021b). Decoding in MTL was lower than in visual cortex, but remained reliably above chance (localizer peak true-class probability: 11.5%, chance: 6.25%; *t*(49) = 22.57, *p* < .001; SI Fig. 6A). Classifiers also generalized to the main task, showing above-chance evidence for presented stimuli during stimulus onset (7.66%, *t*(50) = 13.35, *p* < .001; SI Fig. 6B). Importantly, prior work has demonstrated that sub-second replay can be detected even under substantially reduced classifier accuracy and signal-to-noise conditions, indicating that the comparatively weaker decoding in the MTL does not per se preclude reliable identification of sequential reactivation (e.g., Fig. 5 in Wittkuhn et al. (2021), SI in **?**).

The MTL showed small but consistent sequential replay effects relative to baseline for all sequences, as reflected in a non-zero intercept (*β* = −0.002, *z* = −2.99, *p* = 0.003), and did not appear to differentiate between sequence conditions (main effect condition: *β* = −0.001 for all sequence conditions; see Fig. 5H). Replay in MTL appeared to covary with the strength of visual replay, as evidenced by a mixed-effects model predicting visual replay from MTL replay (*β* = 0.036, *z* = 2.998, *p* = 0.003), indicating cross-region coupling in replay. This coupling was consistent across all conditions (all interaction terms with condition *p >* 0.2). MTL replay did not show any of the modulation of reversal phases, non-local value updates, or relation to structure learning that we observed in visual cortex. Hence, while MTL showed above chance reactivation and correlated with visual replay, it seemed less structured.

To ensure the above results do not reflect inflated statistical baselines, we performed the same analysis on a brain region where we did not expect replay to occur, the somatosensory cortex. No evidence of replay was observed during prolonged ISIs in this region (intercept: *β* = 0.000, SE = 0.001, *z* = 0.096, *p* = 0.923, 95% CI [−0.001, 0.001]; all condition effects *p >* 0.43, coefficients near zero; see Fig. 5H), ruling out the potential issue of an inflated chance baseline that could have affected our results. Both the MTL and the visual ROI significantly differed from this baseline (MTL: *β* = −0.003, *z* = −3.448, *p* = 0.001; visual: *β* = −0.006, *z* = −7.879, *p <* 0.001).

### Replay adapts to changes in reward structure

Our results on Day 1 suggested that replay was more tightly coupled with non-local value learning than structure learning. To assess the sensitivity of replay to structure and change more directly, participants returned for a second session on the following day, during which we manipulated their knowledge of the latent structure and the reward correlations, while keeping all other task aspects identical. To ensure that all participants began Day 2 with the same understanding of the structure learned on Day 1, we explicitly revealed which sequences shared the same reward pairing at the end of Day 1. Before the structure was revealed, participants completed a questionnaire indicating which sequences they believed shared a reward level (i.e., yielded a similar number of coins). Nearly half of the participants (49%) identified the correct set of sequences immediately. After receiving a hint that three paths shared the same reward level, an additional 37% selected the correct paths. The remaining 14% responded incorrectly twice.

On Day 2, participants then began by completing two blocks of the task with the previously revealed reward structure from Day 1. As expected, this led to near-ceiling performance in reversal-locked trials (97%, ± 0.012; Fig. 6B, C, D), and reduced the previously observed difference between semantic and non-semantic generalization (*t*(49) = 1.35, *p* = 0.18), suggesting participants fully relied on the task structure. After block 2, participants were informed that the underlying graph structure would change in the upcoming block. No information about how it would change was provided, such that participants had to discover the new reward structure through experience. The main difference of the new structure was that the A^∗^ path was now coupled with latent reward R2 instead of R1 (Fig. 6A). This meant that the A^∗^ path led to the same outcomes as C, instead of sharing outcomes with A and B as before, and changed the original generalization patterns in two ways: First, some of the generalizations participants had learned on Day 1 were not valid anymore (e.g. A^∗^ ⇒B, A⇒A^∗^, henceforth *No Gen (New)*, dashed brown arrows in Fig. 6A); second, other generalizations that were invalid on Day 1 now became valid (A^∗^⇒C, C⇒A^∗^, henceforth *Gen (New)*, green in Figs 6A, F-H). Our main question was how this change in task structure led to adaptations in behavior and replay. Specifically, we sought to compare generalization and non-generalization conditions that were consistent with the task structure on Day 1 (*Gen (old)* and *No Gen (old)*, respectively) with those consistent with the new task structure (*Gen(new)* and *NoGen(new)*). After completing the fourth block, the new structure was revealed to participants, allowing them to complete the final block with explicit knowledge of the updated reward associations.

**Figure 6:**
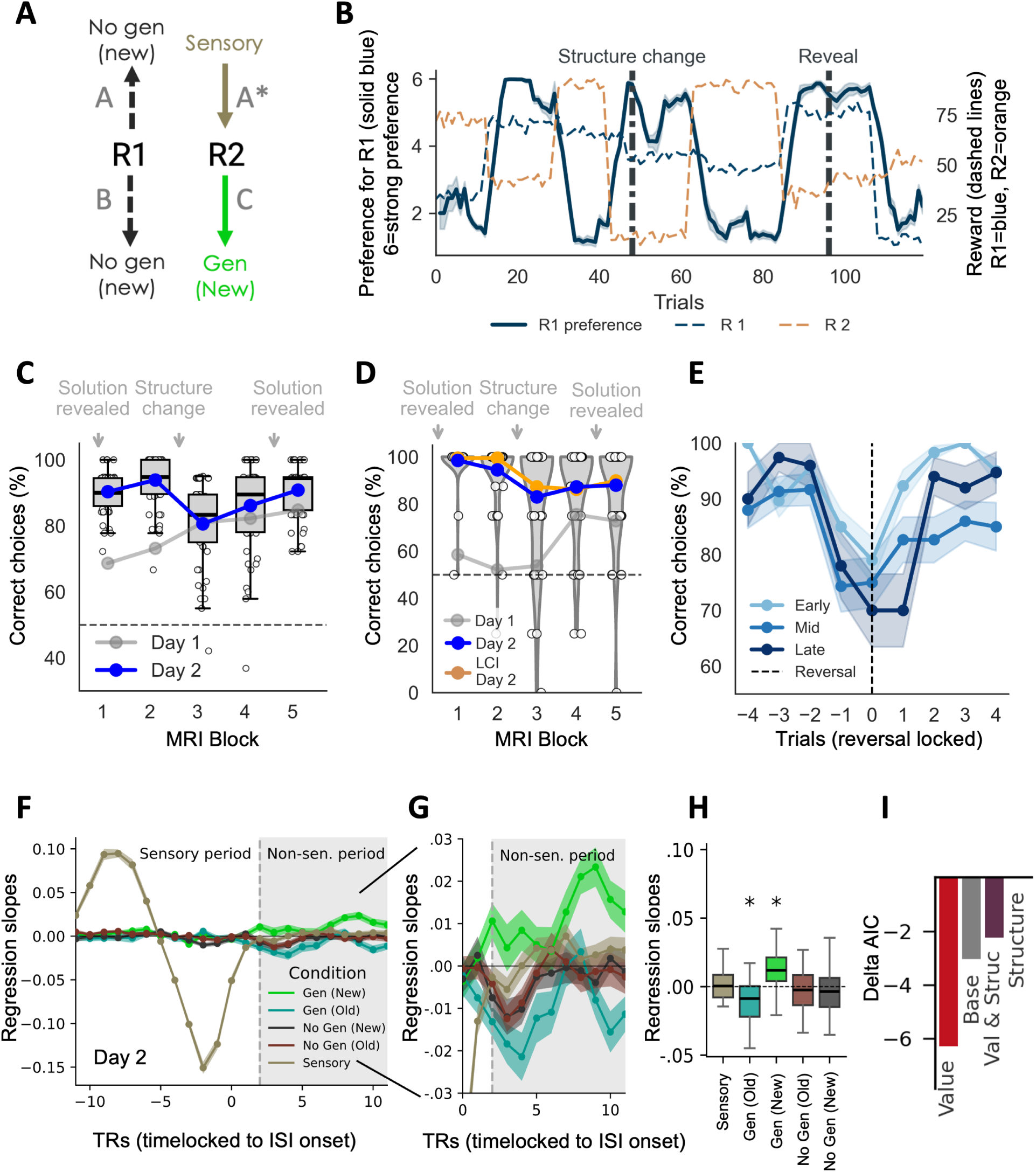
Behavior, LCI, and replay on Day 2. (A) Updated replay and generalization structure following the reassignment of path A* from R1 to R2. Some pre-existing relationships remain unchanged (e.g., generalization across A and B, Gen old; no generalization between A and C, No Gen old), while others reorganize: the former A–A* link becomes invalid (No Gen new) and a new A*–C link emerges (Gen new). (B) Example choice ratings on Day 2, showing preference for R1 (solid blue) alongside the true reward values (dashed lines). The initial phase follows the old structure, followed by a hidden structure change and a later explicit reveal (vertical dashed lines). (C) Accuracy across MRI blocks on Day 2. Participants perform near ceiling early in the session, show a drop after the hidden structure change, and then successfully relearn the updated structure. Day 1 performance (gray) is shown for comparison. (D) Reversal trial accuracy on Day 2. Performance is close to ceiling before the structure change, with a modest drop immediately afterward, indicating rapid adaptation. Day 1 reversal performance (gray) is shown for reference. The LCI model (orange) captures this pattern, reorganizing its latent structure quickly following the change. (E) Reversal recovery on Day 2, time-locked to reversal onset (trial 0). Early reversals occur before the structure change, mid reversals occur after the change but before the reveal, and late reversals occur after the reveal of the new structure. Recovery is fastest during the initial phase; performance after the structure change is slightly reduced but remains above chance, and following the reveal, accuracy returns to near-ceiling within two trials. Note the slight performance dip at trial –1, suggesting that some participants adjusted to emerging reward-level changes before the reversal had fully occurred. (F) Regression slope timecourses on Day 2, showing strong sensory-driven activation and updated replay patterns during the non-sensory period. (G) Zoomed view of the non-sensory period. Sensory and both no-generalization conditions show no reactivation. The old A–B generalization (Gen old) continues to produce robust backward replay, whereas the newly formed A*–C generalization (Gen new) exhibits forward replay. (H) Aggregated regression slopes in the non-sensory period. Both the old (A–B) and new (A*–C) generalization links show significant reactivation; the old link retains backward replay, while the new link exhibits forward replay directionality. (I) Day 2 AIC analysis confirms that trial-by-trial value updating continues to best predict replay reactivation, surpassing a model based on condition alone.

The structure change elicited a drop in accuracy, with some variability across participants (Fig. 6C; paired *t*-test, block 2 vs. block 3: *t*(49) = 7.99, *p <* 0.001). Despite this disruption, participants adapted quickly: performance remained well above chance throughout and improved from 80.6% immediately after the change to 90.8% by the end of Day 2. The recovery following the structure change indicates that the transient drop did not reflect increased difficulty of the new structure but instead participants’ relearning of the reward associations (effect of block after the structure change in the LME: *β* = 0.051, *z* = 5.421, *p <* 0.001). Reversal-trial performance closely mirrored this overall pattern: accuracy dropped from 95% to 83% after the structure change (paired *t*-test, block 2 vs. block 3: *t*(49) = 3.22, *p* = 0.002). Participants remained well above chance and showed a slight improvement across blocks, reaching 88% reversal accuracy by block 5, although this increase was not statistically significant (paired *t*-test, block 3 vs. block 5: *t*(49) = 1.30, *p* = 0.20). This pattern was also evident in the reversal-locked recovery, showing the shallowest drop in the early phase before the structure change, a larger and slower recovery immediately afterward, and a return to early-phase recovery levels once the new structure was revealed, albeit requiring one additional trial after the reversal (Fig. 6E). The LCI model recapitulated the main features of structure relearning on Day 2 (Fig. 6D). When participants encountered an unexpected sequence–reward pairing, the resulting prediction error led the model to infer that a new latent-cause structure was required to account for the observations. This inference initiated the construction of a revised latent structure and drove a gradual reorganization of value estimates around the newly inferred reward-sharing configuration (see SI Fig. 3B). Mirroring human behavior, the model showed a transient drop in predictive accuracy following the contingency change and provided a substantially better fit to Day 2 choices than the temporal-difference models (AICc: 39 vs. 70, 59, and 57 for the TD, 1*α*, and 3*α* models, respectively).

We next analyzed replay in the visual ROI on Day 2. In the first step, we confirmed that results during the early phase of Day 2, where the task structure was unchanged from Day 1, mirrored those observed on Day 1. Indeed, we again found strong sensory-driven sequential activation, and broadly comparable evidence for selective backward reactivation of reward-sharing sequences in visual cortex: the non-semantic generalization model showed significant negative coefficients (*t*(49) = −3.01, *p* = 0.016), and the semantic generalization effect remained negative but did not reach significance (*t*(49) = −1.84, *p* = 0.071; note that only 2 blocks were included resulting in decreased power); the no-generalization sequence showed no reliable effect (*t*(49) = 1.86, *p* = 0.071). Unexpectedly, we observed a significant positive effect of the sensory sequence, consistent with forward sequentiality (*t*(49) = 2.54, *p* = 0.028; see SI Fig. 7).

MTL results in this early phase were also similar to Day 1, showing backward reactivation for all sequence conditions with comparable signal strength: sensory (*t*(49) = −3.26, *p* = 0.008), non-semantic generalization (*t*(49) = −2.08, *p* = 0.043), semantic generalization (*t*(49) = −2.86, *p* = 0.008), and no-generalization (*t*(49) = −2.90, *p* = 0.008). As before, somatosensory cortex showed no significant effects across conditions (all |*t*| ≤ 2.05, all *p* ≥ 0.182).

Analyzing replay following the structure change, we observed that the required relearning elicited a targeted adaptation of replay. While replay of still-valid Gen(old) associations (*A* → *B*) was predominantly backward as before (*t*(49) = −4.90, *p <* 0.001, dark green in Fig. 6F, G, H), we found evidence for above chance *forward* replay that reflected the newly valid generalizations (Gen(new), light green *A*^∗^ → *C*, *C* → *A*^∗^, *t*(49) = 4.91, *p <* 0.001), a finding that held when directly comparing the coefficients of the same condition before vs. after structure change (*t*(49) = 2.28, *p* = 0.027; note that the regression coefficient was part of the no-generalization condition (*A*^∗^ - C) in the early phase, which as we reported above, had a small positive effect already before the structure change; our direct pre vs post comparison accounts for this effect, however). This increase in replay contrasted with the continued absence of replay in the NoGen(old) condition *t*(49) = −1.55, *p* = 0.16, as well as with no evidence of replay for the now-obsolete *A*^∗^–A/B association (NoGen(new); *t*(49) = −1.76, *p* = 0.14, dark brown). In sum, these changes suggest a dynamic adaptation of replay structure that co-occurred with behavioral adaptation to changes in generalization structure.

In the MTL, the structure change did not induce a clear directional flip in replay. All conditions showed evidence of backward sequentiality except the Gen(new), which did not show reliable reactivation in either direction. Specifically, we observed significant negative slopes for the sensory condition (*t*(49) = −3.11, *p* = 0.008), the Gen(old) condition (*t*(49) = −2.46, *p* = 0.029), the No Gen(old) condition (*t*(49) = −2.32, *p* = 0.031), and the No Gen(new) condition (*t*(49) = −3.47, *p* = 0.006), whereas the Gen(new) condition did not reach significance (*t*(49) = −1.11, *p* = 0.272). As before, the somatosensory control region did not reveal significant replay evidence in any condition (all |*t*| ≤ 1.89, all *p* ≥ 0.323).

To ask which factors could explain the replay we again extracted structure learning and value updating metrics from the LCI model for Day 2. Consistent with the previous session, we found no evidence that structure updating predicted replay beyond a baseline model including condition only (AIC = −7871.5 vs. −7874.6; likelihood ratio test: *χ*^2^(2) = 4.956, *p* = 0.292). Unlike on Day 1, reversal periods per se were not a significant predictor of replay (AIC = −7873.0; likelihood ratio test: *χ*^2^(2) = 6.479, *p* = 0.166). Yet, value updating from the LCI model continued to predict replay significantly (AIC = −7877.8; likelihood ratio test: *χ*^2^(2) = 11.259, *p* = 0.024; see Fig. 6I). We did not find a reliable relationship between replay strength and behavioral performance on Day 2, likely due to the limited number of trials before the structure change and the shift in replay direction.

### Medial Temporal Lobe Representations Track Structure Learning and Adaptation

Given that generalisation replay relied on structure knowledge, we asked where this task structure was neurally encoded. We hence tested whether visual and MTL representations encoded the reward structure of the task and whether building these representations was associated with replay. We used cross-validated representational dissimilarity matrices (RDMs) capturing the Mahalanobis distances of fMRI activity patterns between all 16 stimuli for each phase (early, late) of each day. The empirical RDMs were compared then to a model RDM that reflected the expected similarities for brain regions that encoded shared reward contingencies, such that stimuli that led to the same reward (A, A^∗^, B on Day 1) were assigned low dissimilarity, and stimuli with different rewards (C vs. others) high dissimilarity (see Fig. 7A). To ensure specificity, we set up a number of control RDMs that captured higher similarity among items from the same path (e.g. A1 and A2 are more similar than A1 and B2), similarity of items from the same category, and items that appeared at the same sequence position (see Methods for details). We then orthogonalized the shared reward model RDM with respect to all of these alternative model RDMs, such that any effect could be uniquely attributed to learned task structure.

**Figure 7:**
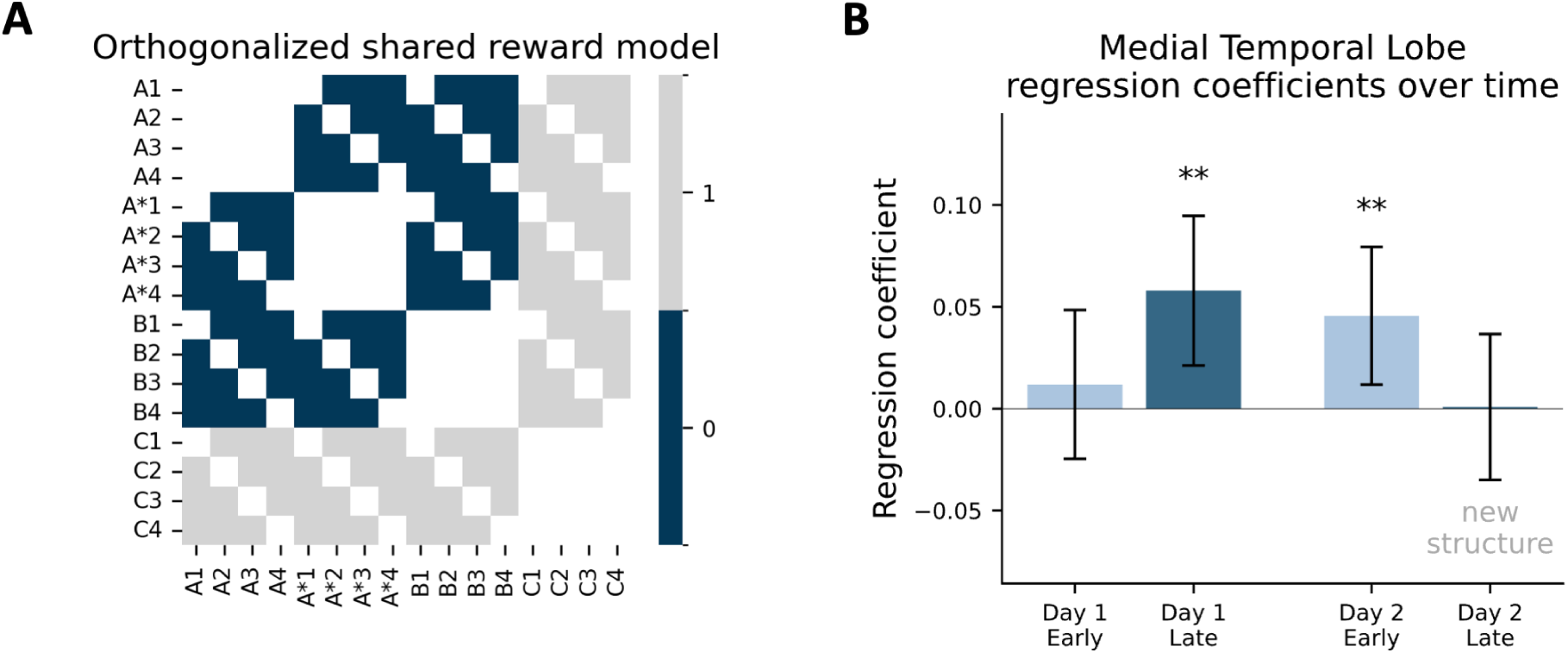
MTL representations track and reorganize according to shared reward structure. (A) Model representational dissimilarity matrix (RDM) encoding shared reward structure. Stimuli from sequences A, A*, and B (leading to R1) are modeled as similar (dark blue), while comparisons to sequence C (leading to R2) are modeled as dissimilar (light gray). The model was orthogonalized against within-sequence similarity, category similarity, and sequence position to isolate reward structure specifically. (B) Regression coefficients showing alignment between empirical MTL representations and the shared reward model across phases. During Day 1, correlations increase from early to late blocks, indicating gradual acquisition of abstract reward structure. On Day 2, alignment is initially maintained but decreases after the structure change when A* switches from R1 to R2, reflecting representational reorganization. Error bars denote 95% confidence intervals of the regression coefficient estimates.

Given previous evidence (Behrens et al., 2018), we first hypothesized that MTL representations would increasingly align with the shared reward model during structure learning on Day 1, remain high when participants began Day 2 with explicit structure knowledge, and decrease after the Day 2 structure change. A mixed-effects model with Day (Day 1, Day 2), phase (early, late), model RDM, and their interactions as predictors revealed the predicted three-way interaction (day × phase × model RDM: *z* = −2.81, *p* = 0.005), indicating that the relationship between MTL representations and the reward structure changed dynamically across learning and adaptation. Follow-up comparisons confirmed the hypothesized pattern. On Day 1, MTL alignment with the shared reward structure increased from early (*β* = −0.001, *z* = −0.05, *p* = 0.96) to late blocks (*β* = 0.054, *z* = 2.86, *p* = 0.004), paralleling behavioral acquisition of the structure (Fig. 7B). On Day 2, alignment was already high in early blocks (*β* = 0.047, *z* = 2.55, *p* = 0.011), reflecting explicit knowledge, but dropped to non-significant levels after the structure change (*β* = −0.006, *z* = −0.29, *p* = 0.77), consistent with representational reorganization during adaptation. Further splitting the empirical RDMs into individual sequence pairs and testing late-early differences revealed small but consistent effects: The only nominally significant effect of similarity decreases on Day 2 was between A and A^∗^, i.e. precisely the pair of arms that was changed (= 0.112, *t*(45) = 2.14, *p* = 0.038). This effect, however, did not survive multiple-comparisons correction (all pairs *p*_FDR_ ≥ 0.40).

The same mixed effect model in the visual ROI showed a significant effect for the structure model RDM (*β* = 1.346, *z* = 4.73, *<* 0.001). However, we found no significant interaction of the model with task phase (early vs. late: (*β* = −0.206, *z* = −0.512, *<* 0.609) or day (*β* = −0.116, *z* = −0.288, *<* 0.773) corresponding to the acquisition during day 1 or the adaptation during day 2. We did not test alignment with the new structure on Day 2 because cross-validated RDMs need to be estimated across blocks. The new structure was valid beginning with block 3, and revealed already after block 4. Hence, cross validation would be based on very little data and would require combining one block with explicit knowledge and one with inferred knowledge, resulting in insufficient power and limited interpretability. We found no significant relationship between changes in representational similarity in MTL and visual or MTL replay activity. Models predicting later similarity change from early replay, and models predicting later replay from representational change, were both non-significant (all *p >* 0.18).

## 3 Discussion

Adaptive decision-making requires the ability to discover latent structure, generalize knowledge to unobserved situations, and flexibly reorganize behavior when that structure changes. Building on computational accounts in which latent-cause inference underlies the discovery of hidden structure from experience (Gershman et al., 2010; Mirea et al., 2024), we apply this framework to model how humans infer latent structure from trial-and-error feedback and use it to flexibly generalize across shared rewards. We also report that the LCI model outperformed classic reinforcement learning models and captured individual differences in structure learning. Neurally, we find evidence for ’non-local’ value propagation as previously reported with MEG (Liu et al., 2021b), albeit we found it localized in visual cortex, rather than in MTL as suggested by (Liu et al., 2021b). Another main novel observation is that replay deeply interacts with latent cause inference, adjusting on a trial by trial basis which experiences are replayed in proportion to the required non-local value updates implemented in a latent cause inference process. Previous work on latent-cause inference has emphasized how experiences separate into distinct latent causes when contingencies change. New latent causes are created when prediction errors signal that the current generative model no longer explains observations, leading to context-specific learning and the preservation of past associations (Gershman et al., 2010). This state-splitting mechanism accounts for phenomena such as renewal after extinction or context-dependent memory retrieval. In contrast, our approach highlights a complementary functioning of latent-cause inference: generalization by assigning different observations to a shared latent cause, similar to Mirea et al. (2024). By mapping distinct experiences onto the same underlying latent cause, the model determines when knowledge learned in one setting can be transferred to another. Thus, rather than proliferating latent causes to explain differences, our account relies on their shared use to integrate information across situations that share an underlying relational structure. This emphasis on generalization links to other representation learning frameworks supporting generalization. The successor representation (SR) (Dayan, 1993; Gershman, 2018; Momennejad et al., 2017; Kahn et al., 2025; Keck et al., 2025) encodes predictive relations among states, forming a cached map of expected future state occupancies that enables rapid reward revaluation but offers limited flexibility when transitions change (but see Wittkuhn et al. (2025) and Sagiv et al. (2025) for evidence linking replay to SR learning and re-learning). The linear reinforcement-learning framework and its default representation (DR) (Piray and Daw, 2021) extend this idea to a stable, policy-independent predictive map that can be reused across changing goals or environments without relearning transition dynamics. Both SR and DR capture structure through prediction rather than inference: by encoding how states unfold through learned transitions, they support generalization by leveraging predictive relations. In contrast, latent-cause inference generalizes by clustering observations under shared hidden causes, allowing the agent to infer when superficially different experiences reflect the same latent structure.

Neurally, we observed two key phenomena that provide insight into how latent structure discovery and neural replay support flexible generalization. First, backward replay in visual cortex selectively reinstated unobserved sequences that shared a latent reward, consistent with a mechanism for non-local value updating (e.g. Liu et al., 2021b) and with the occurrence of learning-related replay in visual cortex (Wittkuhn et al., 2025). Second, representational similarity analyses revealed that multivariate patterns in the MTL gradually aligned with the abstract reward structure, reorganizing when contingencies changed on Day 2.

During the early sensory period, we observed a coactivation of semantically related reward-sharing sequences, exceeding a baseline derived from the localizer and potentially facilitating value transfer between associated items in line with previous results by Wimmer and Shohamy (2012). Following the sensory period after the sensory effect subsided reactivation in visual cortex was selectively expressed for reward-associated sequences, ruling out visually driven or nonspecific reactivation. Replay strength increased across Day 1 as participants discovered the underlying structure, peaked after reward reversals, and predicted performance over a longer behavioral horizon. Different previous accounts disagree about the extent to which replay is hypothesized to support value vs. structure learning Mattar and Daw (2018); Sagiv et al. (2025). In the current study, trial-wise fluctuations in replay were best explained by non-local value updates derived from the LCI model rather than by a structure-learning metric, indicating a tight coupling between inferred latent structure and value propagation. Although negative results should be interpreted with caution — and the current study does not ideally distinguish value from structure learning, which are largely driven by the same reversal events — the lack of evidence for replay involvement in structure learning is striking.

The MTL showed a distinct pattern from visual cortex. While multivariate patterns in the MTL dynamically tracked the reward structure, increasing alignment during Day 1 learning and reorganizing after the Day 2 structure change, MTL replay itself was broad and lacked the selectivity observed in visual areas. This contrasts with previous human replay research utilizing MEG (e.g. Wimmer et al., 2023; Liu et al., 2019; Nour et al., 2021), where we observe different signatures in cortical versus medial temporal regions. Previous simulations confirm that replay can be detected despite reduced SNR (Wittkuhn and Schuck, 2021), and MRI studies have detected replay in the hippocampus (Schuck and Niv, 2019), further validating our MTL findings. Moreover, correlation between visual cortex and MTL replay and the selective presence of reactivation in MTL alongside the absence of replay in a somatosensory control region indicate that the broad MTL replay reflects genuine, albeit less targeted, reactivation. Two interpretations, not mutually exclusive, could explain this regional difference. First, classification in MTL was constrained by modest decoding accuracy, potentially limiting the granularity with which sequence-specific replay can be resolved. Second, MTL replay may reflect more global sampling, providing a structural scaffold over which selective cortical replay operates. This interpretation aligns with Kurth-Nelson et al. (2016), who suggested that MEG-detected replay is driven by sensory regions reflecting MTL-orchestrated reactivation. Under this view, the MTL maintains the abstract task structure and drives the selective replay we observe in cortex.

The structural reversal on Day 2 further revealed how replay tracks the evolving task structure. While backward replay along previously valid generalization links persisted, newly valid links exhibited forward replay. Notably, value updating from our LCI model remained the strongest predictor of replay during relearning. Whether this directional shift reflects replay passively following an already updated representation or actively participating in structural relearning remains an open question.

Together, these findings indicate that latent-structure discovery and neural replay jointly support flexible generalization. Latent causes organize experience into shared reward sources, enabling rapid inference, while replay propagates value across this inferred structure to support non-local generalization. Two key questions remain. First, our reversal manipulation closely coupled structural inference with non-local value updating, making it difficult to isolate their respective contributions. Future work should experimentally dissociate changes in latent structure from changes in value, and track how replay direction evolves as new structure is acquired versus merely exploited. Second, the broad and relatively non-selective replay observed in the MTL remains only partly understood. Clarifying whether this pattern reflects methodological limits on decoding or a coordination signal that scaffolds more selective cortical replay will be essential for understanding how replay and latent structure interact to enable rapid, flexible behavior beyond direct experience.

## 4 Methods

### 4.1 Participants

Fifty-two neurotypical participants (mean age = 27.4 years, SD = 5.6; 25 females) were recruited from the participant database of the Max-Planck Institute for Human Cognitive and Brain Sciences, Leipzig. All participants had no counter indications for MRI scanning, no history of psychiatric conditions, normal or corrected-to-normal vision and provided written informed consent before participation. The study was approved by the ethics committee at the University of Leipzig (002/22-ek). One participant discontinued during the localizer task on day 1 and was excluded entirely. Another participant did not complete day 2 due to illness, and was therefore only included in day 1 analysis. Finally, 3 participants had incomplete scanning blocks during day 1 due to technical malfunctions. In these cases we use all available data, i.e. only excluded blocks were necessary. This resulted in 51 participants for day 1 analyses and 50 participants for day 2 analyses. Participants received €12 per hour ( €96 total for 8 hours across two 4-hour sessions) plus a performance-based bonus up to €34, calculated from points collected during the main tasks and accuracy in localizer tasks.

### 4.2 Experimental Design

The experiment consisted of four fMRI sessions split across two consecutive days. Each day began with a localizer session for classifier training (sessions 1 and 3; 70 minutes), followed by the main task (session 2 and 4; 65 minutes). Both days included a break of approximately 20 minutes between sessions.

#### 4.2.1 Sessions 1 and 3 - localizer

Session 1 (Day 1 Localizer): Participants viewed 17 stimuli in a pseudorandom order while performing a 1-back task. Stimuli consisted of 1 reward image (treasure chest) and 16 unique 750ms video clips of everyday objects, including animals (giraffe, zebra), vehicles (sports car, generic car), buildings (modern house, old house), and food items (raspberries, grapes). Videos were selected to create clear categorical groupings, relevant for the main task, while maintaining visual distinctiveness for classification. Stimuli were presented using PsychoPy (version 2022.2.4) and projected onto a screen visible via a mirror mounted on the head coil. Each stimulus was presented 34 times across six blocks with balanced transitions (579 trials in total). The mean inter stimulus interval was 3s sampled from a truncated exponential distribution ranging from 2 to 7s. On occasional probe trials (30% of trials), participants had to identify the stimulus shown immediately before the probe by a left or right button press. Three types of probe trials were included to ensure attentiveness: (1) the same stimulus and a foil, (2) a different exemplar of the same object category (e.g., a different zebra) and a foil, and (3) two foils, requiring a special button press (down) when neither matched the previously seen object. The maximum response time was 2 seconds. This localizer session provided unbiased training data for the multivariate classifiers. The two localizer sessions across Days 1 and 2 were identical in design. However, participants differed in their knowledge of the task. During the first localizer on Day 1, participants were naive to both the upcoming main task and the latent reward structure. On Day 2, participants again completed the same localizer, but now with full knowledge of the latent structure and the associations among stimuli, although the structure knowledge was not required to perform the localizer task.

#### 4.2.2 Sessions 2 and 4 - main task

The main task involved four sequential “bandits” (A, A*, B, and C), each consisting of a fixed sequence of four 750 ms video clips that culminated in a reward outcome. On Day 1, sequences A, A*, and B were linked to a shared latent reward (R1), whereas sequence C was associated with an independent reward (R2). Rewards were matched in accumulated value across all trials. Observed rewards ranged between 0 and 100 and were stochastically sampled from the true mean of R1 or R2 on each trial with a small standard deviation (SD = 2). Throughout the session, true reward means were organized into stable phases punctuated by six reversals (15–20 trials apart). During reversals, mean rewards changed abruptly, typically passing through 1–2 transitional values before stabilizing at the new level. Each participant was assigned to one of four reward schedules (different schedules each day) generated by counterbalancing (i) the order of reward changes and (ii) which of the two latent reward timecourses was assigned to R1 vs. R2. This ensured that group-level effects could be attributed to learning rather than idiosyncratic reward trajectories. Although reversals of R1 and R2 sometimes co-occurred (see Fig. 2A/B), reward time-courses were designed to also include phases in which one reward changed while the other remained stable, as well as intervals of gradual drift and abrupt shifts (average correlation between R1 and R2 time courses: r = -0.18). Assignment of video identities to sequence positions was randomized across schedules. To provide an additional scaffold for structure learning, sequences A and A* also shared semantic category membership at corresponding positions (e.g., both showing animals at position 1, houses at position 2, etc.; the category ordering varied across participants), thereby creating a “semantic generalization” condition (shared reward and shared category), in addition to “non-semantic generalization” (shared reward only). For example, A: giraffe → modern house → raspberries → sports car; A*: zebra → old house → grapes → generic car. The first day concluded with explicit instructions about the latent reward structure.

Day 1 main task (Session 2) comprised 120 trials arranged in 5 blocks, with each sequence shown six times per block. Each trial consisted of a sequential presentation of four videos (750 ms each) followed by a reward image (750 ms) depicting coins and overlaid with the associated numerical outcome. After observing each sequence and its outcome, participants made preference judgments between two sequence elements (4 s response window). Participants were instructed to choose the option they believed was associated with the more valuable reward at that moment. Confidence was indicated on a 6-point scale, with extreme ratings (1 or 6) reflecting high confidence and values closer to the center indicating greater uncertainty. 100 such judgments were collected over the session, always matching positions within the sequence; positions 2 and 3 were probed more frequently, with probe frequencies of 6, 45, 30, and 9 trials for positions 1–4, respectively. Ten trials involved options that originated from the same reward source and therefore lacked a correct choice under the initial structure (e.g., choices between A and A*, which only acquired different values after the structure change). On these trials, participants used the slider less decisively, showing more center-position ratings rather than strong preferences at the extremes, as confirmed by a linear mixed-effects model predicting ratings based on whether the probed options came from shared versus separate reward sources. Five trials following each reward reversal served as critical tests of latent structure knowledge, requiring participants to generalize value from an observed sequence to an unobserved one. For example, after a reversal, the first four trials might only present sequences A and C, while the preference judgments probe A* and B. On the fifth trial, either A* or B is shown and the remaining unobserved option is probed. Successfully performing these trials therefore requires inferring the relative value of unobserved sequences based on the most recent observed outcome. Each day participants solved 6 reversals which were distributed across the 5 blocks, with blocks 2 or 4 containing 2 reversals.

The reward schedules consisted of stable phases with small fluctuations, gradual drifts, and occasional abrupt changes, creating instances in which reversal choices could be solved based on reward memory. These trials still required adapting reward preferences (i.e., switching from preferring R1 to R2 or vice versa) but could be guided by previously observed outcomes if the drifting reward source had not yet changed substantially. For example, when the shared reward R1 drifted gradually while R2 abruptly dropped from high to low, observing the new R2 outcome together with one R1 branch provided sufficient information to infer the values of the remaining (unprobed) R1 branches. In such cases, only minor value updating through generalization was needed, allowing participants to rely on prior experience. When focusing on trials that required updating reward preferences but could not be guided by previously observed outcomes (e.g., when both rewards changed simultaneously), learning improvements from early (Blocks 1–2) to late (Blocks 4–5) stages were comparable to those observed across all reversal trials. The session concluded with explicit feedback about the latent reward structure.

On Day 2 main task (Session 4), participants first received a reminder of the latent reward structure and completed a short behavioral training block outside the scanner to practice reward-based generalization. Inside the scanner, Blocks 1–2 then assessed performance with full structure knowledge. After Block 2, participants were informed that the underlying reward structure would change, but were not told how. In Blocks 3–4, the reward contingency of A* switched from R1 to R2, requiring participants to update their generalization strategy. Before Block 5, the new latent structure was explicitly revealed. This manipulation required participants to abandon previously learned semantic associations and base generalization exclusively on the new reward contingencies. Inter-stimulus intervals (ISIs) were drawn from a truncated exponential distribution (mean = 3 s, range 2–7 s). On a subset of trials, the ISI following reward was prolonged by an additional 15 s (average 18 s total) before the preference judgment. These prolonged ISIs occurred eight times per block (40 per session) and were concentrated around reward reversals to provide periods without sensory input during which neural replay was expected to support non-local value updating.

### 4.3 Behavioral Analysis

Performance was quantified using multiple metrics. Overall accuracy was defined as the percentage of trials in which participants selected the option leading to the higher reward. To assess generalization after reversals, we analyzed the accuracy of post-reversal choices involving bandits whose outcomes had not yet been observed since the reversal (i.e., choices requiring generalization from observed to unobserved sequences). Learning dynamics were further characterized by computing trial-wise accuracy time-locked to reward reversals (Fig. 6 E).

To examine the influence of category information, we analyzed how the category of the observed sequence affected subsequent choices on reversal trials. Specifically, we compared generalization performance for within-category generalization (e.g., A→A* or A*→A) versus across-category generalization (e.g., B→A*/A or A*/A→B) to test whether semantic similarity between observed and generalized sequences facilitated structure discovery (Fig. 2 E).

Statistical analyses were conducted using linear mixed-effects models implemented in Python 3.11.7 using the package statsmodels (version 0.14.4). Models included a random intercept for each participant and tested the fixed effects of interest. Statistical tests were done using scipy (version 1.12.0). Reversal performance analyses (Fig. 2C, 6D) were corrected for multiple comparisons using false discovery rate correction.

### 4.4 Latent Cause Inference Model

To formalize how participants discover and update latent task structure, we fitted a latent cause inference (LCI) model (e.g. Gershman et al., 2010; Pisupati et al., 2024; Mirea et al., 2024; Berwian et al., 2025) that assigned observations to latent causes using a Chinese Restaurant Process (CRP) prior (Aldous, 2006). The model assumed that the observed sequence–reward pair (*s_t_, r_t_*) on a given trial was generated by an unobserved latent cause with time varying properties, *z_t_*. Each cause was characterized by (i) a categorical distribution over the four sequences and (ii) a reward expectation. The model then used approximate Bayesian inference to infer which cause had generated the observation, and updated the causes correspondingly. If participants made a choice between two stimuli, without observing any reward, the model retrieved the associated reward expectations given the shown stimuli. The model involved four mechanisms: (1) computation of a prior of latent causes, (2) computation of likelihoods of observed stimuli and reward given each latent cause, (3) sequential Bayesian inference of the posterior using particle filtering, (4) value estimation and decision making. These steps are described in detail below. Model parameters were fitted for each subject by maximum likelihood estimation as described below.

#### CRP prior

The LCI model maintained a prior over latent causes *z_i_* given the previous latent causes in all preceding trials. The main idea of the CRP prior we used here is that the probabilities of existing causes were proportional to how often they had occurred in previous trials, while the probability to generate new causes was governed by a concentration parameter *α* and decreased over time:

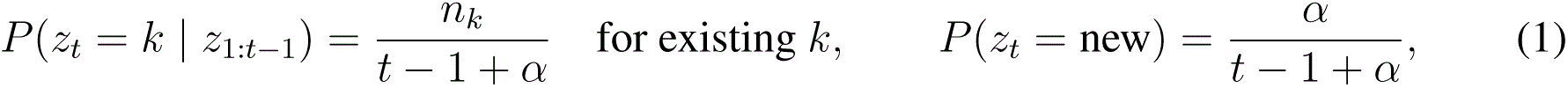

We denote the latent causes on trial *t* by *z_t_*. Let *k* index an existing cause, with *n_k_* the number of previous assignments to cause *k*. Thus, the CRP prior favors reusing existing causes in proportion to how often they have been inferred before, while allowing occasional creation of new causes controlled by the concentration parameter *α*.

#### Likelihoods

##### Categorical sequences

Each latent cause maintains a Dirichlet distribution that describes how probable each of the four possible sequences is given each cause. Note that, for simplicity, we do not model the semantic similarity between these sequences, and instead treat them as four distinct categorical observations.At the beginning of the experiment we assume a flat Dirichlet with pseudo-counts (1, 1, 1, 1). Let *n_k,s_* denote the number of times sequence *s* has been assigned to cause *k*. A category-increment parameter *η* controls how strongly each observation updates the categorical distribution:

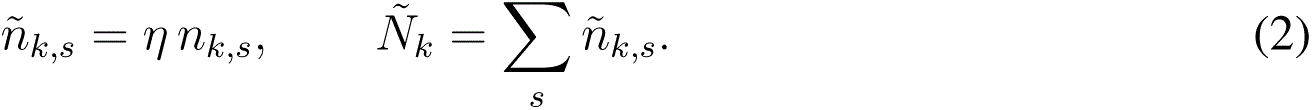

The categorical probability for existing causes is then

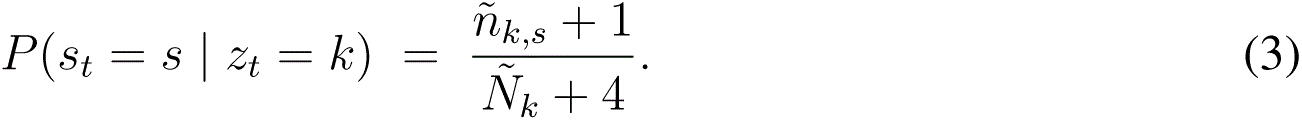

For a newly created latent cause, we use the uniform categorical prior *P* (*s_t_*= *s* | *z_t_* = new) = 1*/*4.

##### Rewards

Reward likelihood is a uniform–Gaussian mixture:

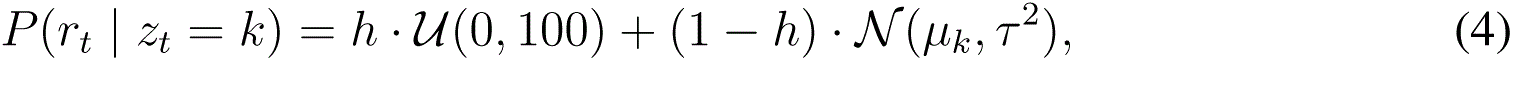

with hazard rate *h* (e.g., Nassar et al., 2010). The uniform component allows abrupt reward changes to be incorporated without necessitating creation of new latent causes, preserving the shared structure needed for value generalization. In the online update used here, *µ_k_*is set to the most recent reward assigned to cause *k* and a fixed *τ* ^2^ of 5 is used for existing causes; for a new cause we use *µ* = 50 and a broader variance of 100.

#### Computing the posterior and particle filtering

Because exact Bayesian inference of the posterior is non-tractable, we approximate inference using a particle filtering approach. The above process of computing the prior and likelihoods was repeated for each of 100 particles, on each trial (Sanborn et al., 2010). This process involved the following steps for every particle.

1. Compute the CRP prior over existing causes plus a potential new cause.
2. For each candidate cause, compute the likelihood combining categorical and reward probabilities *P* (*s_t_* | *z_t_* = *k*) *P* (*r_t_* | *z_t_*= *k*) (for potential new latent causes using uniform sequence and broad reward likelihood)
3. Branch the particle for each candidate cause and assign the weight of each candidate by prior×likelihood. This creates for each particle multiple candidates (one per existing cause and an additional for a new cause).
4. Update the categorical and reward estimate of each candidate based on the current observation.
5. Normalize branch weights across the full candidate pool.
6. Resample back to 100 particles; weights are reset to uniform after resampling.

#### Calculating value estimates and value-based decision-making

After each observation, the particle filter expands and resamples the particle set. Each particle *i* ∈ {1*, … , M* } maintains a set of currently instantiated latent causes *K_i_*, each with an associated reward mean *µ_ik_* and categorical counts over sequences. The categorical posterior-predictive *P_i_*(*s* | *k*) for each cause is defined as in the Categorical sequences section.

For a queried sequence *s*, each particle forms a mixture over its own latent causes by normalizing these categorical predictive probabilities:

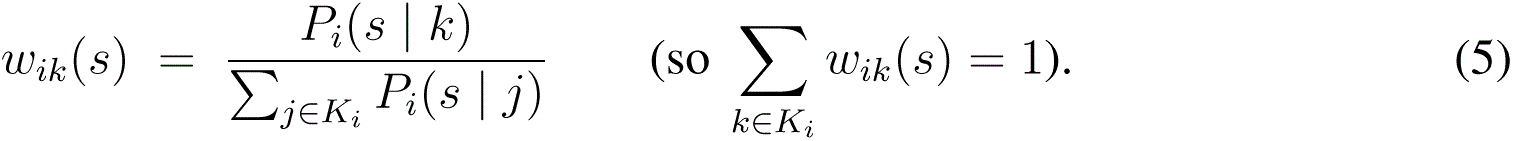

The particle-level value of sequence *s* is then the expectation of the reward means under these mixture weights:

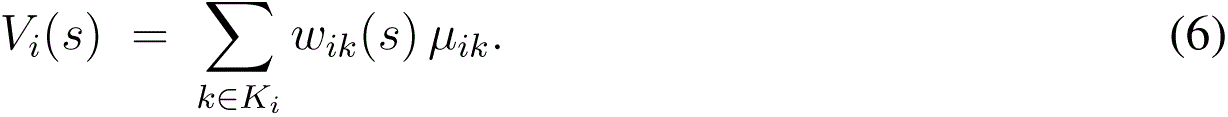

Because particles are resampled and assigned equal weight after every trial, the model’s predicted value is the simple average across particles:

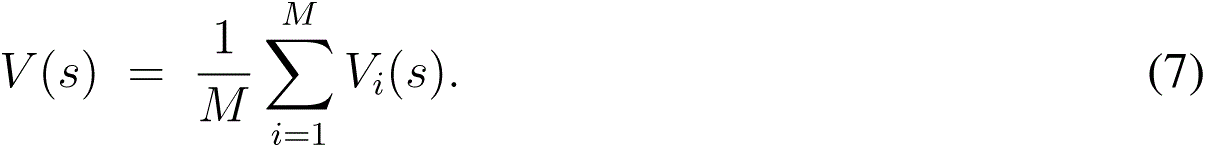

Intuitively, each particle represents a distinct hypothesis about how observations cluster into latent causes; *V* (*s*) averages these hypotheses to obtain the predicted expected reward for each sequence which builds the basis for subsequent decision-making.

To produce binary choices between two sequences *s*_1_ and *s*_2_, the model first computes the value of each sequence as described above, by averaging each particle’s reward expectation weighted by its latent-cause mixture probabilities. These values *V* (*s*_1_) and *V* (*s*_2_) are then transformed into choice probabilities using a softmax function with inverse-temperature parameter *θ*:

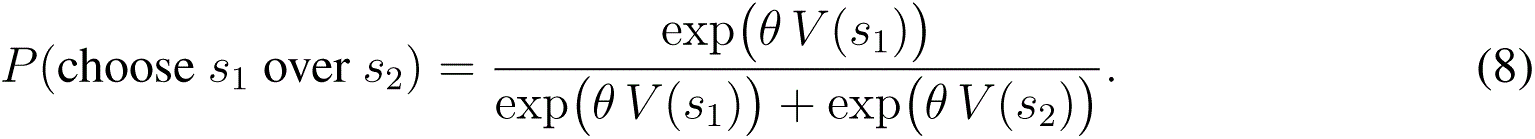

#### Parameter estimation and model comparison

Participant-specific parameters were estimated for each day separately by maximizing the likelihood of observed choices using PyBADS (Acerbi and Ma, 2017). The LCI model had the following 4 free parameters and bounds:

- *α* ∈ [0, 1]: CRP concentration (new latent cause control)
- *θ* ∈ [0, 1]: softmax inverse-temperature (choice stochasticity)
- *h* ∈ [0, 1]: hazard rate (mixture of uniform and gaussian reward distribution)
- *η* ∈ [1, 10]: category increment (evidence weight per sequence observation)

We used a structured multi-start (3 starts per parameter; 3^4^ = 81 runs) and selected the best log-likelihood. Model comparison used the corrected Akaike Information Criterion (AICc):

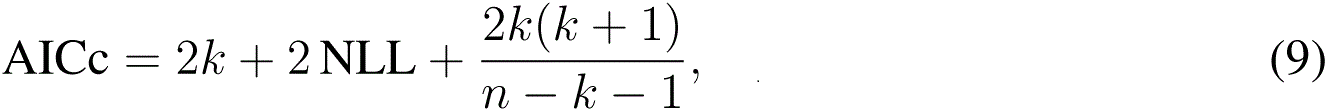

computed per participant.

To elucidate the potential computational functions of replay, we derived two trialwise metrics from the fitted latent cause inference (LCI) model. The first metric captured nonlocal value updating, quantifying the extent to which observing a reward led to changes in the inferred values of sequences that were not directly experienced on that trial. This measure was computed on a sequence- and trial-specific level, allowing us to link replay-related neural activity to the model’s propagation of reward information across related task elements (Fig. 5E). The second metric reflected structure learning, tracking how the model inferred latent task structure over time. For each trial, we obtained the probability distribution over latent causes for every sequence and quantified their pairwise similarity by computing symmetric Kullback–Leibler (KL) divergences between these distributions. Smaller KL divergences indicated increasing overlap in latent-cause representations, reflecting convergence toward a shared structural understanding independent of reward (Fig. 5F). To match the trialwise dynamics of nonlocal value change, we additionally computed the trialwise change in the KL divergence metric, providing a dynamic measure of structural updating across trials. To align model dynamics with the experimental manipulation of task structure on Day 2, the model was initialized with two latent causes, each assigned to the corresponding sequences with a pseudocount of 5. After the first two blocks, the categorical likelihoods and priors were broadened by a fixed decay parameter of 0.8 to simulate increasing uncertainty, and following block 4, the latent-cause assignments were updated to reflect the new post-change reward structure. The resulting trialwise estimates of both nonlocal value updating and structure learning were used as predictors in mixed-effects models linking behavior and replay (Fig. 5G).

#### Alternative temporal difference learning models

To dissociate flexible latent-structure learning from simpler accounts, we compared the LCI model against temporal-difference (TD) models with and without hard-coded generalization. All TD variants track a separate value *V* (*s*) for each sequence *s* ∈ {*A, B, A*^∗^*, C*} and update the value of the experienced sequence *s_t_*on trial *t* via a standard delta rule,

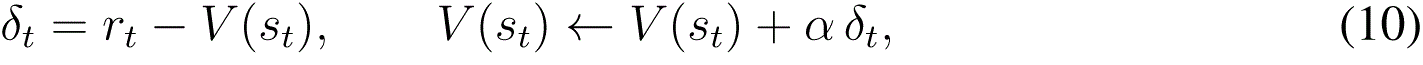

where *r_t_* is the observed reward and *α* a learning rate. In the *no-generalization* baseline, unvisited sequences are never updated, preventing generalization and requiring values to be relearned independently after reversals. In the *hard-coded generalization* variants, the same prediction error *δ_t_* is also applied to predefined related sequences according to the task’s latent structure (e.g., between *A* and *A*^∗^, and between *A/A*^∗^ and *B*). One variant uses a single learning rate *α* for both direct and generalized updates, enforcing fixed-strength generalization from the outset. A second variant uses three separate learning rates *α*_exp_ for direct experience, *α*_semantic_ for generalization between *A* and *A*^∗^, and *α*_non−semantic_ for generalization between *A/A*^∗^ and *B*, allowing graded influence along different generalization types. All TD models translate values into choices using the same softmax choice rule and inverse-temperature *θ* used by the LCI model. Critically, because their generalization structure is prespecified and static, these models cannot discover latent structure, nor flexibly adapt their structure when reward contingencies change (e.g., on Day 2).

### 4.5 MRI

#### 4.5.1 MRI data acquisition and preprocessing

MRI data were acquired using a 32-channel head coil on a 3T Siemens Magnetom Prisma Fit system. Functional MRI (fMRI) scans were collected in a transverse orientation using T2*-weighted gradient-echo echo-planar imaging (GE-EPI) with multiband acceleration, sensitive to BOLD contrast. This sequence achieves high temporal resolution while maintaining good spatial coverage and resolution. We acquired 72 transverse slices with a thickness of 2 mm and an in-plane resolution of 2 × 2 mm. Imaging parameters included a multiband acceleration factor of four, a repetition time (TR) of 1.5 seconds, and an echo time (TE) of 24.2 ms. The field of view (FoV) was 204 mm in both read and phase directions, and slices were collected in an interleaved order. Fat saturation was applied, and the flip angle was set to 70°. The first five volumes of each block were discarded to allow for scanner equilibration. Additionally, each day a high-resolution T1-weighted anatomical scan (1 × 1 × 1 mm) was acquired for spatial normalization and coregistration. To correct for susceptibility-induced distortions, we collected field maps using spin-echo EPI sequences with opposing phase encoding polarities (anterior-posterior and posterior-anterior), which were used for distortion correction. All fMRI preprocessing steps were performed using fMRIPrep.

Results included in this manuscript come from preprocessing performed using *fMRIPrep* 23.1.4 (Esteban et al. (2019); Esteban et al. (2018); RRID:SCR 016216), which is based on *Nipype* 1.8.6 (Gorgolewski et al. (2011); Gorgolewski et al. (2018); RRID:SCR 002502).

##### Preprocessing of B0 inhomogeneity mappings

A total of 6 fieldmaps were found available within the input BIDS structure for this particular subject. A *B0*-nonuniformity map (or *fieldmap*) was estimated based on two (or more) echo-planar imaging (EPI) references with `topup` (Andersson et al. (2003); FSL None).

##### Anatomical data preprocessing

A total of 2 T1-weighted (T1w) images were found within the input BIDS dataset. All of them were corrected for intensity non-uniformity (INU) with `N4BiasFieldCorrection` (Tustison et al., 2010), distributed with ANTs (version unknown) (Avants et al., 2008, RRID:SCR 004757). The T1w-reference was then skull-stripped with a *Nipype* implementation of the `antsBrainExtraction.sh` workflow (from ANTs), using OASIS30ANTs as target template. Brain tissue segmentation of cerebrospinal fluid (CSF), white-matter (WM) and gray-matter (GM) was performed on the brain-extracted T1w using `fast` (FSL (version unknown), RRID:SCR 002823, Zhang et al., 2001). An anatomical T1w-reference map was computed after registration of 2 T1w images (after INU-correction) using `mri robust template` (FreeSurfer 7.3.2, Reuter et al., 2010). Brain surfaces were reconstructed using `recon-all` (FreeSurfer 7.3.2, RRID:SCR 001847, Dale et al., 1999a), and the brain mask estimated previously was refined with a custom variation of the method to reconcile ANTs-derived and FreeSurfer-derived segmentations of the cortical gray-matter of Mindboggle (RRID:SCR 002438, Klein et al., 2017). Volume-based spatial normalization to two standard spaces (MNI152NLin6Asym, MNI152NLin2009cAsym) was performed through nonlinear registration with `antsRegistration` (ANTs (version unknown)), using brain-extracted versions of both T1w reference and the T1w template. The following templates were were selected for spatial normalization and accessed with *TemplateFlow* (23.0.0, Ciric et al., 2022): *FSL’s MNI ICBM 152 non-linear 6th Generation Asymmetric Average Brain Stereotaxic Registration Model* [Evans et al. (2012), RRID:SCR 002823; TemplateFlow ID: MNI152NLin6Asym], *ICBM 152 Nonlinear Asymmetrical template version 2009c* [Fonov et al. (2009), RRID:SCR 008796; TemplateFlow ID: MNI152NLin2009cAsym].

##### Functional data preprocessing

For each of the 30 BOLD runs found per subject (across all tasks and sessions), the following preprocessing was performed. First, a reference volume and its skull-stripped version were generated using a custom methodology of *fMRIPrep*. Head-motion parameters with respect to the BOLD reference (transformation matrices, and six corresponding rotation and translation parameters) are estimated before any spatiotemporal filtering using `mcflirt` (FSL, Jenkinson et al., 2002). The estimated *fieldmap* was then aligned with rigid-registration to the target EPI (echo-planar imaging) reference run. The field coefficients were mapped on to the reference EPI using the transform. BOLD runs were slice-time corrected to 0.699s (0.5 of slice acquisition range 0s-1.4s) using `3dTshift` from AFNI (Cox and Hyde, 1997, RRID:SCR 005927). The BOLD reference was then co-registered to the T1w reference using `bbregister` (FreeSurfer) which implements boundary-based registration (Greve and Fischl, 2009). Co-registration was configured with six degrees of freedom. Several confounding time-series were calculated based on the *pre-processed BOLD*: framewise displacement (FD), DVARS and three region-wise global signals. FD was computed using two formulations following Power (absolute sum of relative motions, Power et al. (2014)) and Jenkinson (relative root mean square displacement between affines, Jenkinson et al. (2002)). FD and DVARS are calculated for each functional run, both using their implementations in *Nipype* (following the definitions by Power et al., 2014). The three global signals are extracted within the CSF, the WM, and the whole-brain masks. Additionally, a set of physiological regressors were extracted to allow for component-based noise correction (*CompCor*, Behzadi et al., 2007). Principal components are estimated after high-pass filtering the *preprocessed BOLD* time-series (using a discrete cosine filter with 128s cut-off) for the two *CompCor* variants: temporal (tCompCor) and anatomical (aCompCor). tCompCor components are then calculated from the top 2% variable voxels within the brain mask. For aCompCor, three probabilistic masks (CSF, WM and combined CSF+WM) are generated in anatomical space. The implementation differs from that of Behzadi et al. in that instead of eroding the masks by 2 pixels on BOLD space, a mask of pixels that likely contain a volume fraction of GM is subtracted from the aCompCor masks. This mask is obtained by dilating a GM mask extracted from the FreeSurfer’s *aseg* segmentation, and it ensures components are not extracted from voxels containing a minimal fraction of GM. Finally, these masks are resampled into BOLD space and binarized by thresholding at 0.99 (as in the original implementation). Components are also calcuated separately within the WM and CSF masks. For each CompCor decomposition, the *k* components with the largest singular values are retained, such that the retained components’ time series are sufficient to explain 50 percent of variance across the nuisance mask (CSF, WM, combined, or temporal). The remaining components are dropped from consideration. The head-motion estimates calculated in the correction step were also placed within the corresponding confounds file. The confound time series derived from head motion estimates and global signals were expanded with the inclusion of temporal derivatives and quadratic terms for each (Satterthwaite et al., 2013). Frames that exceeded a threshold of 0.5 mm FD or 1.5 standardized DVARS were annotated as motion outliers. Additional nuisance timeseries are calculated by means of principal components analysis of the signal found within a thin band (*crown*) of voxels around the edge of the brain, as proposed by (Patriat et al., 2017). The BOLD time-series were resampled into standard space, generating a *preprocessed BOLD run in MNI152NLin6Asym space*. First, a reference volume and its skull-stripped version were generated using a custom methodology of *fMRIPrep*. The BOLD time-series were resampled onto the following surfaces (FreeSurfer reconstruction nomenclature): *fsnative*, *fsaverage*. All resamplings can be performed with *a single interpolation step* by composing all the pertinent transformations (i.e. head-motion transform matrices, susceptibility distortion correction when available, and co-registrations to anatomical and output spaces). Gridded (volumetric) resamplings were performed using `antsApplyTransforms` (ANTs), configured with Lanczos interpolation to minimize the smoothing effects of other kernels (Lanczos, 1964). Non-gridded (surface) resamplings were performed using `mri vol2surf` (FreeSurfer).

##### Functional data preprocessing

For each of the 30 BOLD runs found per subject (across all tasks and sessions), the following preprocessing was performed. First, a reference volume and its skull-stripped version were generated using a custom methodology of *fMRIPrep*. Head-motion parameters with respect to the BOLD reference (transformation matrices, and six corresponding rotation and translation parameters) are estimated before any spatiotemporal filtering using `mcflirt` (FSL, Jenkinson et al., 2002). The estimated *fieldmap* was then aligned with rigid-registration to the target EPI (echo-planar imaging) reference run. The field coefficients were mapped on to the reference EPI using the transform. BOLD runs were slice-time corrected to 0.7s (0.5 of slice acquisition range 0s-1.4s) using `3dTshift` from AFNI (Cox and Hyde, 1997, RRID:SCR 005927). The BOLD reference was then co-registered to the T1w reference using `bbregister` (FreeSurfer) which implements boundary-based registration (Greve and Fischl, 2009). Co-registration was configured with six degrees of freedom. Several confounding time-series were calculated based on the *pre-processed BOLD*: framewise displacement (FD), DVARS and three region-wise global signals. FD was computed using two formulations following Power (absolute sum of relative motions, Power et al. (2014)) and Jenkinson (relative root mean square displacement between affines, Jenkinson et al. (2002)). FD and DVARS are calculated for each functional run, both using their implementations in *Nipype* (following the definitions by Power et al., 2014). The three global signals are extracted within the CSF, the WM, and the whole-brain masks. Additionally, a set of physiological regressors were extracted to allow for component-based noise correction (*CompCor*, Behzadi et al., 2007). Principal components are estimated after high-pass filtering the *preprocessed BOLD* time-series (using a discrete cosine filter with 128s cut-off) for the two *CompCor* variants: temporal (tCompCor) and anatomical (aCompCor). tCompCor components are then calculated from the top 2% variable voxels within the brain mask. For aCompCor, three probabilistic masks (CSF, WM and combined CSF+WM) are generated in anatomical space. The implementation differs from that of Behzadi et al. in that instead of eroding the masks by 2 pixels on BOLD space, a mask of pixels that likely contain a volume fraction of GM is subtracted from the aCompCor masks. This mask is obtained by dilating a GM mask extracted from the FreeSurfer’s *aseg* segmentation, and it ensures components are not extracted from voxels containing a minimal fraction of GM. Finally, these masks are resampled into BOLD space and binarized by thresholding at 0.99 (as in the original implementation). Components are also calculated separately within the WM and CSF masks. For each CompCor decomposition, the *k* components with the largest singular values are retained, such that the retained components’ time series are sufficient to explain 50 percent of variance across the nuisance mask (CSF, WM, combined, or temporal). The remaining components are dropped from consideration. The head-motion estimates calculated in the correction step were also placed within the corresponding confounds file. The confound time series derived from head motion estimates and global signals were expanded with the inclusion of temporal derivatives and quadratic terms for each (Satterthwaite et al., 2013). Frames that exceeded a threshold of 0.5 mm FD or 1.5 standardized DVARS were annotated as motion outliers. Additional nuisance timeseries are calculated by means of principal components analysis of the signal found within a thin band (*crown*) of voxels around the edge of the brain, as proposed by (Patriat et al., 2017). The BOLD time-series were resampled into standard space, generating a *preprocessed BOLD run in MNI152NLin6Asym space*. First, a reference volume and its skull-stripped version were generated using a custom methodology of *fMRIPrep*. The BOLD time-series were resampled onto the following surfaces (FreeSurfer reconstruction nomenclature): *fsnative*, *fsaverage*. All resamplings can be performed with *a single interpolation step* by composing all the pertinent transformations (i.e. head-motion transform matrices, susceptibility distortion correction when available, and co-registrations to anatomical and output spaces). Gridded (volumetric) resamplings were performed using `antsApplyTransforms` (ANTs), configured with Lanczos interpolation to minimize the smoothing effects of other kernels (Lanczos, 1964). Non-gridded (surface) resamplings were performed using `mri vol2surf` (FreeSurfer).

Many internal operations of *fMRIPrep* use *Nilearn* 0.10.1 (Abraham et al., 2014, RRID:SCR 001362), mostly within the functional processing workflow. For more details of the pipeline, see the section corresponding to workflows in *fMRIPrep*’s documentation.

#### 4.5.2 Copyright Waiver

The above boilerplate text was automatically generated by fMRIPrep with the express intention that users should copy and paste this text into their manuscripts *unchanged*. It is released under the CC0 license.

#### 4.5.3 Multivariate Pattern Analysis for Replay Detection

We trained multivariate pattern classifiers to decode individual stimulus representations from fMRI activity patterns, following established methods for detecting fast sequential reactivation (Schuck and Niv, 2019; Wittkuhn and Schuck, 2021; Wittkuhn et al., 2025). Classifiers were optimized and trained exclusively on task-independent localizer data (Sessions 1 and 3), where all 16 video stimuli were presented with balanced frequencies and transitions. Following Wittkuhn and Schuck (2021), participant-specific anatomical ROIs were defined on the cortical surface using FreeSurfer’s automated parcellation of individual T1-weighted reference images obtained during fMRIPrep preprocessing (Dale et al., 1999b). The occipitotemporal ROI included the cuneus (1005/2005), lateral occipital sulcus (1011/2011), pericalcarine gyrus (1021/2021), superior parietal lobule (1029/2029), lingual gyrus (1013/2013), inferior parietal lobule (1008/2008), fusiform gyrus (1007/2007), inferior temporal gyrus (1009/2009), parahippocampal gyrus (1016/2016), and middle temporal gyrus (1015/2015). The medial temporal lobe ROI comprised the hippocampus (17/53), parahippocampal gyrus (1016/2016), and entorhinal cortex (1006/2006). As a control ROI, we included primary somatosensory cortex, defined as the postcentral gyrus (1022/2022). Within each anatomical ROI, voxel selection and classifier hyperparameters were optimized using a nested cross-validation procedure. Voxel selection was based on two criteria: activation strength and reliability. First, we quantified activation strength as the mean BOLD signal intensity across the entire time course, excluding voxels with poor signal quality or signal dropout.

Second, voxel reliability was assessed using a split-half correlation approach (Tarhan and Konkle, 2020). Beta estimates were derived from a run-wise first-level GLM implemented in Nilearn (SPM canonical HRF; cosine drift model; high-pass filter = 1/128 Hz; voxel-wise standardization; version 0.10.3). For each run, the design matrix included onsets and durations for all 16 video stimuli (modeled as 0.75 s events) and, when present, a 2 s *probe* condition. Nuisance regressors were added per run and comprised the global signal, framewise displacement, six motion parameters and their temporal derivatives (12 total), and six anatomical CompCor components (aCompCor00–05). Voxel reliability was then computed by correlating beta response profiles between odd (runs 1/3/5) and even (runs 2/4/6) localizer runs. For each voxel, condition-wise beta maps were averaged within odd and even sets to yield paired response profiles across all regressors of interest. Pearson correlations between these profiles produced voxel-wise reliability maps and allowed the selection of the reliable subset of voxels for subsequent analyses.

Activation and reliability thresholds were determined within the nested cross-validation and applied iteratively to identify the combination yielding optimal classification performance. In visual cortex, best performance was achieved when selecting the 50% most active and 70% most reliable voxels, using training data combined across both localizer sessions (Sessions 1 and 3). In contrast, for the MTL, optimal performance was obtained with more restrictive thresholds, selecting the 30% most active and 30% most reliable voxels. Notably, training the MTL classifier across both sessions decreased performance relative to session-specific training. Thus, for subsequent main-task analyses and replay detection, the MTL classifier was trained on Session 1 only. We implemented one-versus-rest logistic regression classifiers with L2 regularization, chosen for their efficiency with high-dimensional data and probabilistic outputs necessary for replay detection. Inside the nested cross-validation, we additionally optimized spatial smoothing, where best cross-validated classification was obtained with 4mm FWHM spatial smoothing for all regions. We further applied temporal smoothing: For each stimulus, we extracted classifier training data from the TR closest to 5s post-stimulus onset and the preceding TR in the visual cortex. For the MTL, we obtained best classification smoothing over the pre- and succeeding TR (i.e. averaging over TRs 4-6), maximizing signal-to-noise ratio while capturing peak BOLD response. To assess classifier validity, we first examined potential confusions between stimuli. Classifiers assigned only marginally higher probabilities to stimuli within the same category compared to other stimuli, indicating little to no category-level bias (see confusion matrix, Fig. 4 A). We then validated performance during sequence observation in the main task, where classifiers successfully decoded all 16 stimuli with distinct probability peaks aligned to stimulus presentation times (Fig. 4 C). One classifier corresponding to a house stimulus was excluded from subsequent replay analyses. During reward presentation (the treasure chest), which was not included in sequence modeling, this classifier showed increased decoding probabilities, suggesting a reward-evoked modulation of its time course that could otherwise be confounded with replay-related reactivation.

#### 4.5.4 Sequential Replay Detection

We applied the sequential order dynamics analysis (SODA) from Wittkuhn and Schuck (2021); Wittkuhn et al. (2025) to detect sequential reactivation of stimulus representations during extended rest periods. SODA leverages the temporal dynamics of overlapping hemodynamic responses: when a neural sequence is rapidly replayed, each element evokes a hemodynamic response offset in time. Due to the sluggish nature of the BOLD signal, these overlapping responses create characteristic temporal patterns in classifier probability time courses. The key insight behind SODA is that probabilistic classifiers applied to overlapping BOLD activity produce systematic fluctuations that preserve information about sequential order. To evaluate the proposed sequentiality for every TR the proposed ordering is regressed onto the probability estimates. If stimuli are replayed in the order S_1_ → S_2_ → S_3_ → S_4_, the resulting classifier probabilities will show ordered dynamics: initially, earlier items (e.g., S_1_) will have higher probabilities than later items (e.g., S_4_), creating a forward-ordered pattern. As the hemodynamic responses evolve, this pattern reverses, with later items showing higher probabilities, a backward-ordered pattern.

To test whether such ordering exists, we inserted eight extended 18-second intervals per block (40 per session), during which participants viewed only a fixation cross. These 18-second pauses occurred immediately following reward presentation, i.e. before decision prompts and concentrated around reward reversals when non-local updating is critical. We then applied trained classifiers for all 16 stimulus categories to every TR within these intervals, such as to observe spontaneous reactivation without external input during these time windows. To test for sequential reactivation, we categorized potential replay based on relationships to the just-observed sequence into four possible sequential (re)activation events, as described in the main text: Sensory sequential activation, replay that reflects semantic generalization, replay that reflects non-semantic generalization and replay of sequences to which reward information could not be generalized.

For each category and TR, we quantified sequentiality by regressing sequence position (1–4) against classifier probabilities for the four stimuli comprising that sequence. The resulting regression coefficient reflects both the strength and direction of sequential ordering. To distinguish stimulus-driven responses from post-sensory reactivation, we modelled the full sensory response and defined the transition point following the full forward and backward phases of the sequentiality signal as the boundary between “sensory” and “non-sensory” phases. At this point, classifier probabilities for the just presented (sensory) stimuli returned to chance level, and regression coefficients of the sequentiality metric are back at 0. All subsequent effects in the non-sensory phase therefore cannot be attributed to ongoing sensory input and are interpreted as reflecting internally generated sequence reactivation. All analyses focused on full four-item sequences rather than pair-wise transitions, differing from alternative approaches focusing on pairwise transitions (Liu et al., 2021a).

#### 4.5.5 Representational similarity analysis

We employed representational similarity analysis (RSA) (Kriegeskorte et al., 2008) to examine how neural representations of the 16 task stimuli reorganized to reflect the underlying reward structure during learning and adaptation. RSA quantifies the similarity structure of neural patterns by computing pairwise distances between all conditions, resulting in a representational dissimilarity matrix (RDM). By comparing empirical RDMs to theoretical model RDMs, we can test specific hypotheses about how information is organized. We constructed a binary model RDM encoding the shared reward structure, where stimuli leading to the same reward show low dissimilarity while those with different rewards show high dissimilarity. Specifically, the model RDM contained:

- Low dissimilarity (0) for stimulus pairs sharing rewards: A-B, A-A*, B-A*, and similarities within sequences
- High dissimilarity (1) for stimulus pairs with different rewards: all comparisons between {A, B, A*} and C

To account for potential confounds, we constructed control RDMs encoding alternative organizational principles: within-sequence similarity (e.g., elements within the A sequence), categorical similarity, and sequential position (all first elements across sequences). The shared reward model was orthogonalized with respect to these control models to ensure effects were specifically driven by reward structure rather than perceptual or sequential factors.

To obtain the empirical RDM, we estimated stimulus-evoked BOLD responses using mass-univariate GLMs implemented in Nilearn. Each of the 5 functional runs per main task session was modeled independently using a standard GLM approach with canonical SPM hemodynamic response function, AR(1) noise model, 4 mm FWHM spatial smoothing, and high-pass filtering (128 s cutoff).

Design matrices included 18 stimulus regressors (one per video stimulus, plus choice and sequence-onset events) convolved with the HRF, and 14 nuisance regressors (global signal, frame-wise displacement, 6 motion parameters, 6 anatomical CompCor components). Stimuli were modeled as 750 ms events at presentation onset; choice periods used actual reaction times as durations. For each stimulus, we computed t-statistics contrasting stimulus-specific activation against baseline, yielding 16 t-maps per run. Following the GLM, we computed empirical RDMs based on the multivariate noise-normalized cross-validated Mahalanobis distance (Walther et al., 2016) as implemented in the PyRSA toolbox (https://github.com/Charestlab/pyrsa) (Nili et al., 2014). We estimated the empirical RDM for each experimental phase (early blocks (1&2) and late blocks (4&5) of each day), resulting in a total of 4 empirical RDMs (early and late for each day). This approach allowed to balance stable parameter estimates using cross-validation, while still resolving the temporal dynamics unfolding over the course of the task, capturing the initial learning phase during day 1 followed by the relearning on the second day.

Based on prior literature, we focused on the MTL as a region often associated with representing abstract task structure. ROIs were defined using the same Freesurfer approach described in the replay section, combined with the activation and reliability thresholds. To test our hypothesis of dynamic structure learning, we employed a linear mixed-effects model testing the three-way interaction:

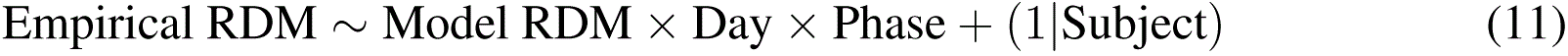

This interaction tests whether representational alignment with the task structure increases from early to late on Day 1 (structure learning) and decreases from early to late on Day 2 (adaptation to structure change)

## Data availability

The data and code supporting the findings of this study will be made available upon publication.

## Acknowledgments

FMR was supported by the Max Planck School of Cognition. PD was supported by the Max Planck Society and the Humboldt Foundation. C.F.D.’s research is supported by the Max Planck Society and the Kavli Foundation. NWS was supported by the European Research Council (ERC Starting Grant REPLAY-852669) as well as the Federal Ministry of Education and Research (BMBF) and the Free Hanseatic City of Hamburg under the Excellence Strategy of the Federal Goverment and the Laender. The views expressed in this article are those of the authors and do not reflect the opinions of the funders.

