## Supplementary material for "Neural replay is connected to latent cause inference and supports fast generalization": SI_Figure_1_to_7

### Supplementary Information for Neural replay is connected to latent cause inference and supports fast generalization

January 16, 2026

#### Remaining behavioral schedules

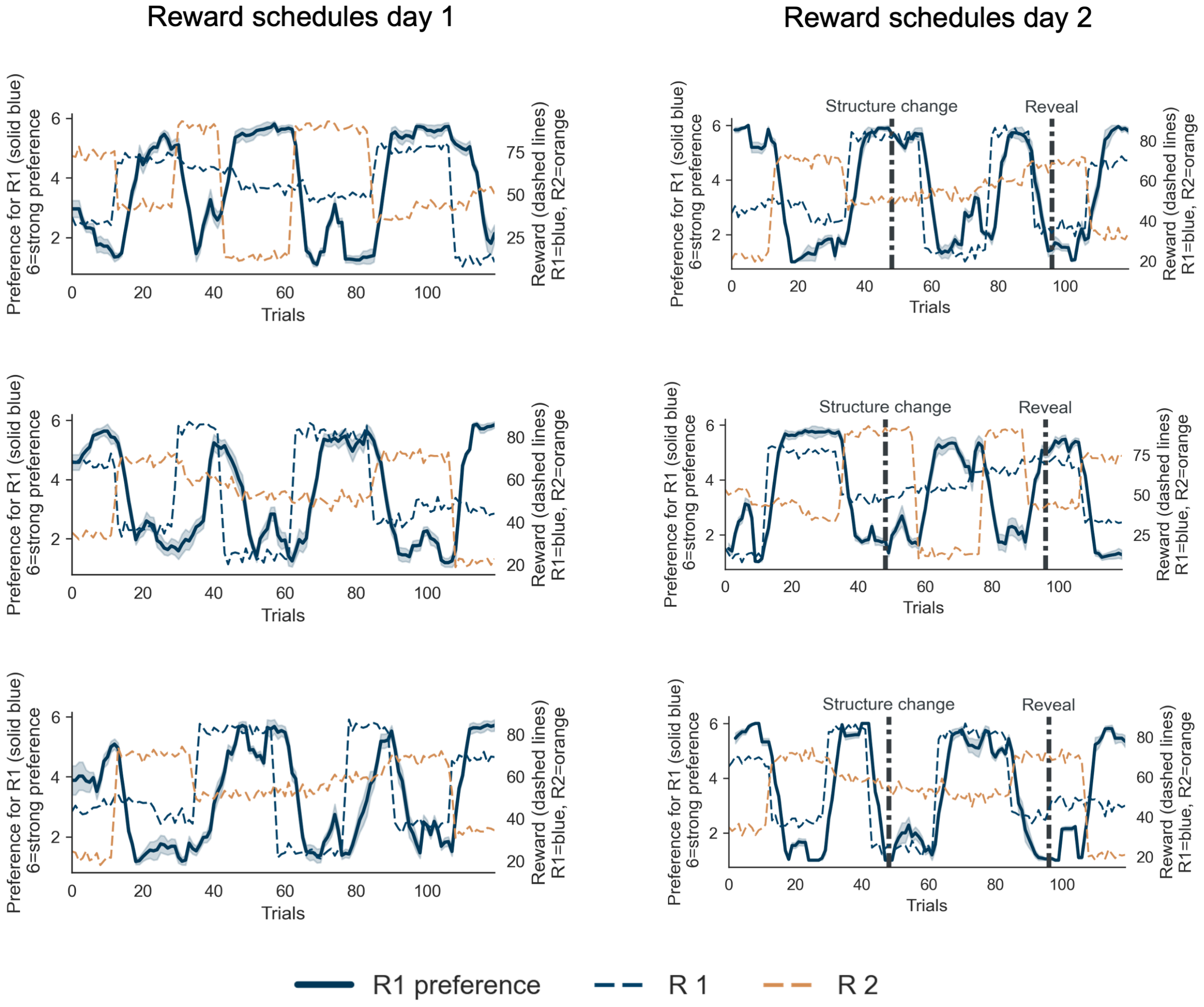


Figure 1: **Choice ratings and reward schedules across days.** Left column: the three remaining reward schedules for Day 1, showing trial-wise choice preference (solid blue) alongside the true reward means for R1 and R2 (dashed lines). Right column: the corresponding schedules for Day 2, including the hidden structure change and subsequent explicit reveal.

#### Latent cause inference model

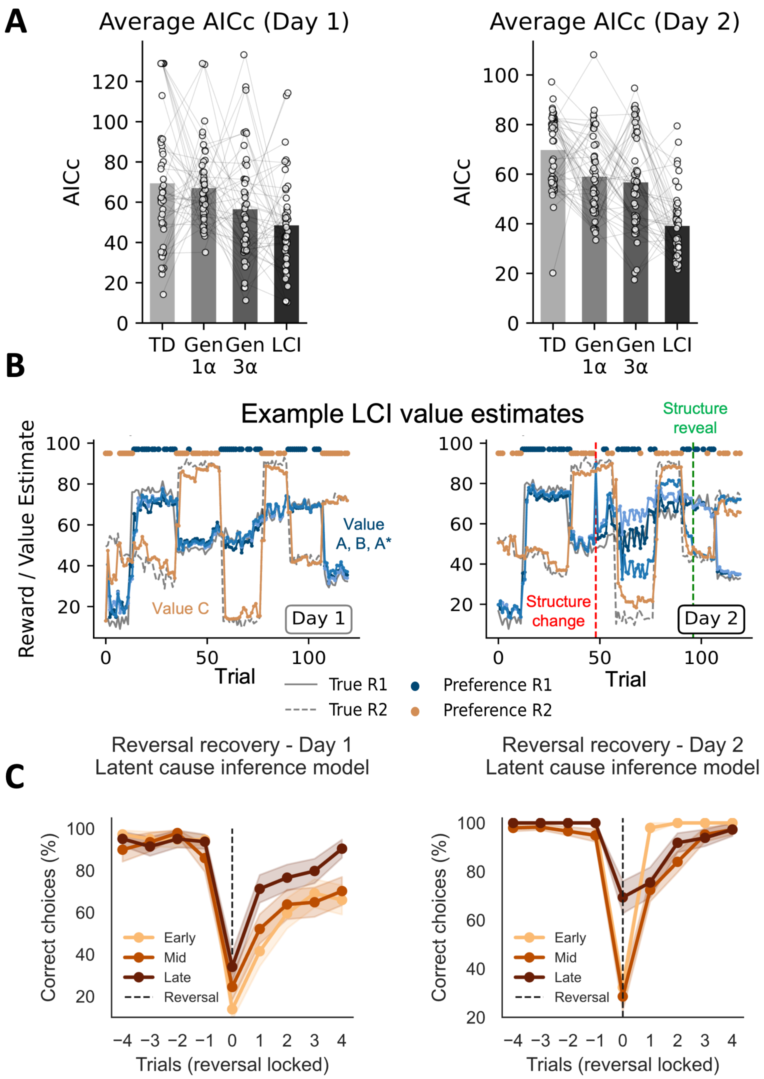


Figure 2: **LCI model comparison, latent-cause value trajectories, and reversal adaptation** (A) AICc scores for all models on Day 1 and Day 2. Across both days, the latent cause inference (LCI) model provides the best fit to participants' choices, outperforming both non-generalizing temporal-difference (TD) learning and TD models with hard-coded generalization (Gen 1, Gen 3). Dots show individual participants; bars show group means. (B) Example trial-wise value estimates generated by the LCI model. On Day 1, the model converges onto the shared reward structure linking A, A*, and B, while treating C as independent. On Day 2, value estimates reorganize when A* switches reward sources (red line), and stabilize once the new structure is explicitly revealed (green line). Grey traces display the true underlying reward means; colored traces show the model-inferred values. (C) Reversal behavior of the LCI model. The model captures the sharp drop in accuracy at reversals (trial 0) and the subsequent recovery. Recovery accelerates from early to late blocks on Day 1 as structure knowledge improves, and remains robust on Day 2 despite the hidden structure change.

#### Localizer task session 1 and 3 and stimuli

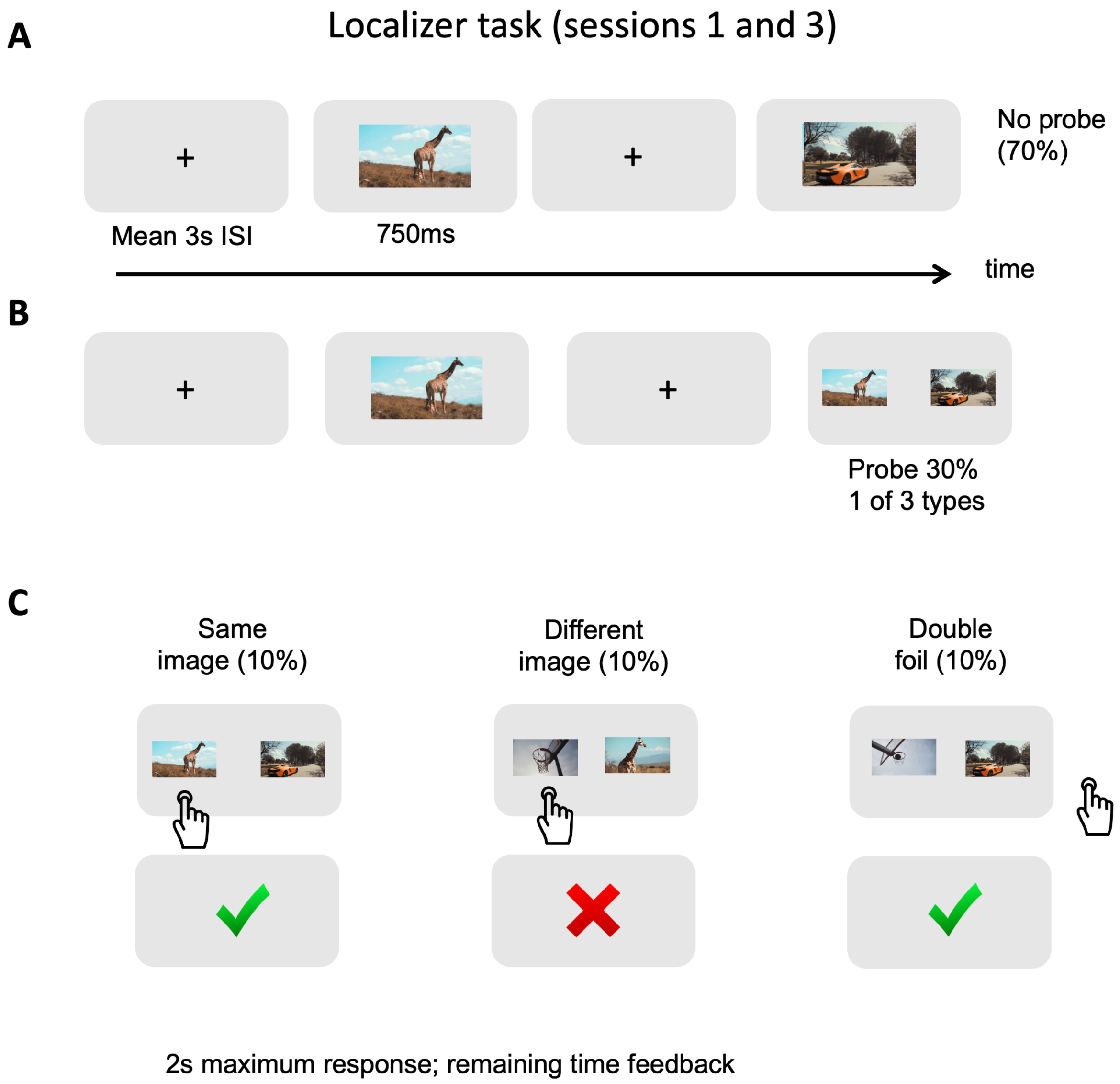


Figure 3: **Localizer task (sessions 1 and 3).** (**A**) Trial structure during non-probe trials (70%). Each image was displayed for 750 ms, separated by a mean inter-stimulus interval of 3 s (range: 2–7 s). (**B**) On probe trials (30%), one of three probe types followed the stimulus. (**C**) Probe types required indicating whether: the same exemplar reappeared (same image), a different exemplar from the same category appeared (different image), or neither image matched the preceding stimulus (double foil; correct down-arrow response). Participants had up to 2 s to respond, after which accuracy feedback was shown for the remainder of the interval.

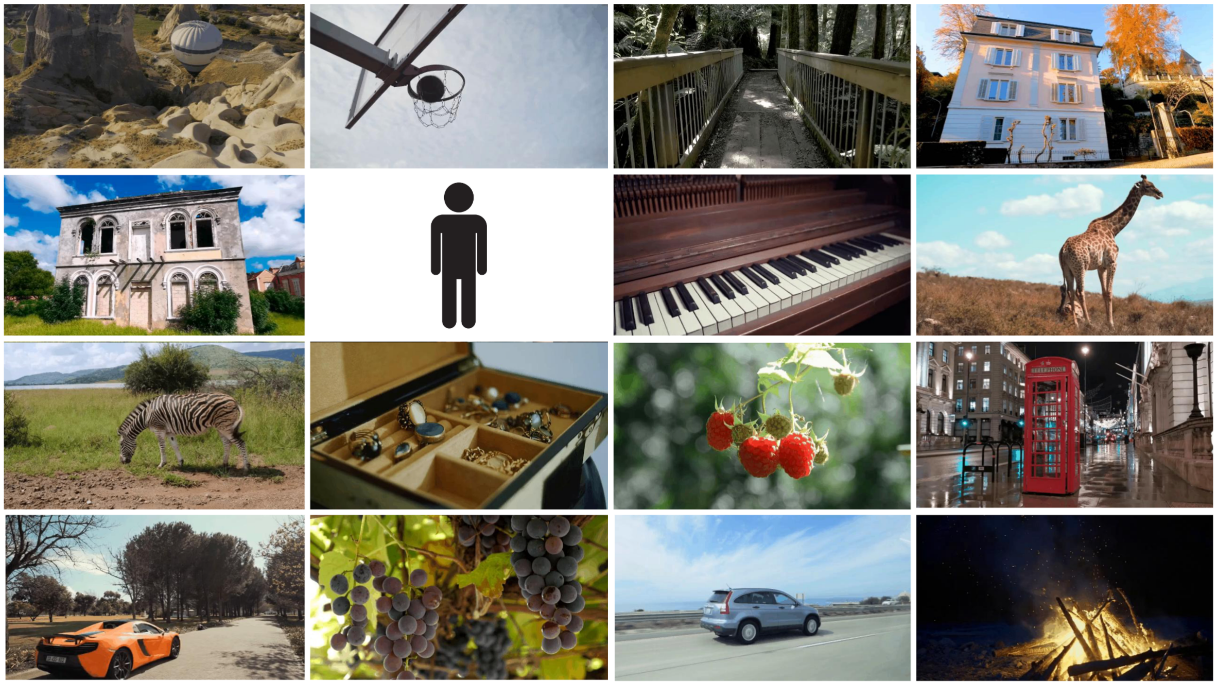

Figure 4: **Stimulus set.** Example stimuli used in the experiment. The shared-category items consisted of two exemplars each from four categories: cars, houses, animals, and fruits. All original images were sourced from the free stock image database Pexels. One stimulus depicting a human was replaced with a schematic human icon to comply with privacy and identifiability restrictions.

#### Decoding performance

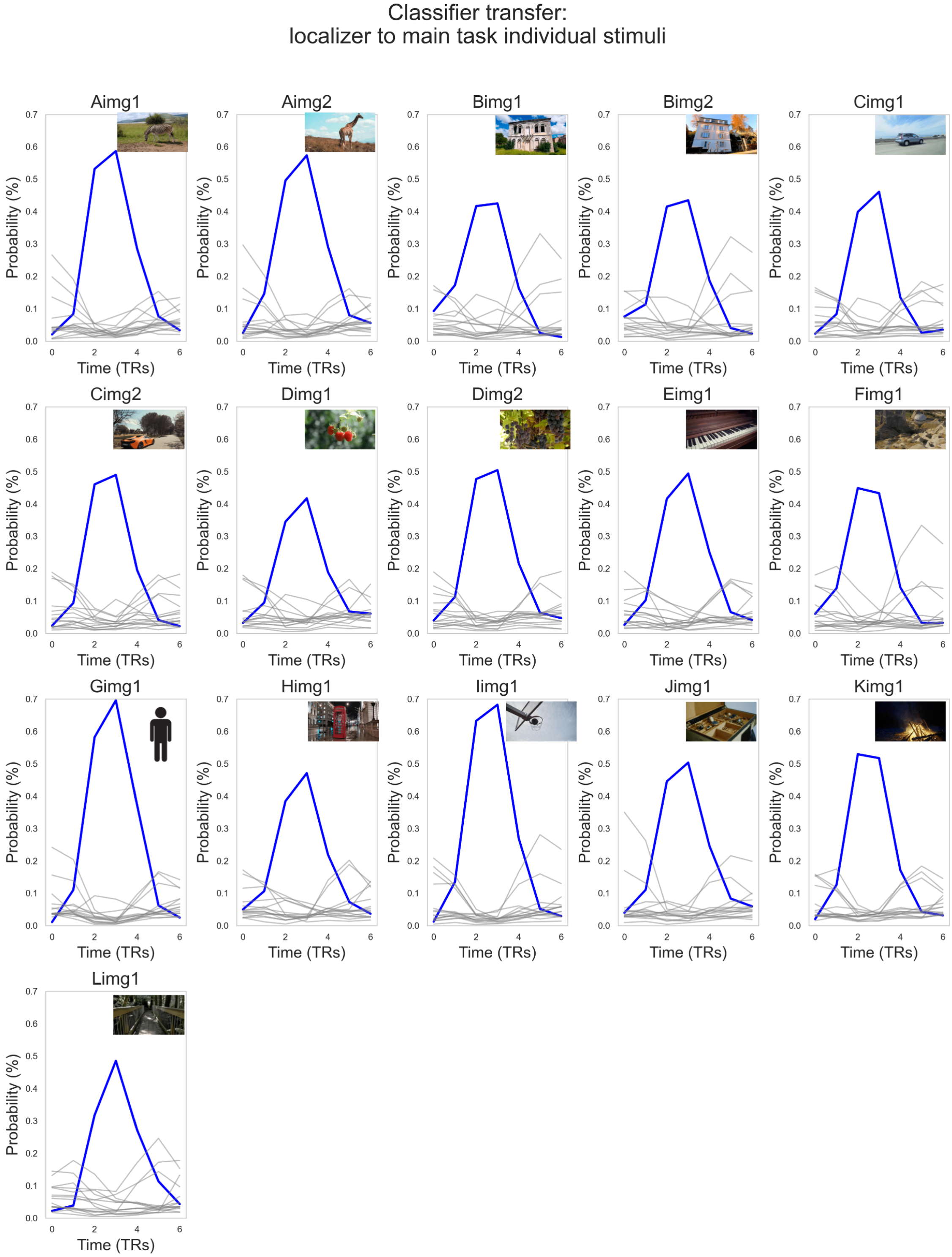


Figure 5: **Classifier transfer from localizer to main task.** Time-resolved classifier outputs (in TRs) for each of the 16 stimuli, decoded from main task data using the localizer-trained classifier. For each stimulus, the corresponding class (blue) exhibits a strong, stimulus-locked probability increase, while nonmatching classes (grey) remain low. This pattern demonstrates robust stimulus-specific decoding and successful transfer of the classifier from the localizer scans to the main task.

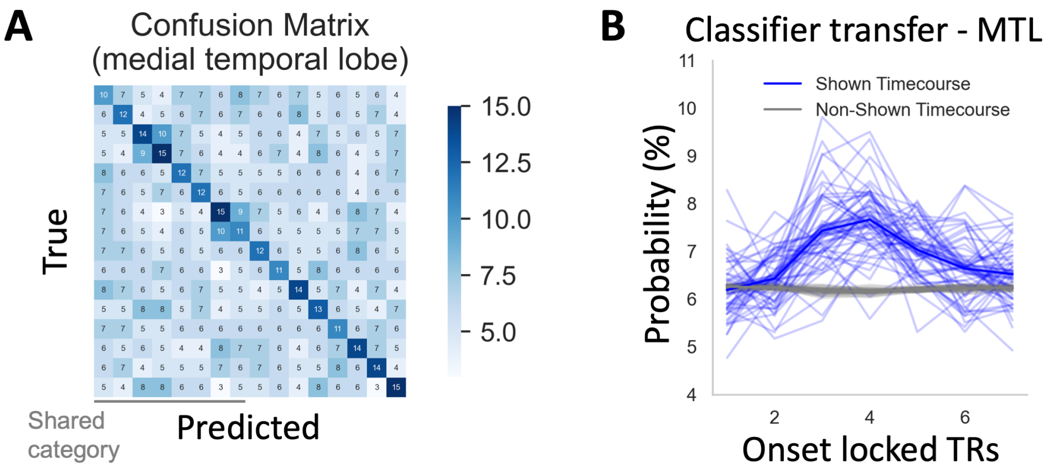


Figure 6: **Decoding for medial temporal lobe.** (A) Confusion matrix for all 16 stimuli decoded within the medial temporal lobe ROI. A clear diagonal indicates successful stimulus-level decoding despite overall lower decoding accuracy in this region. Off-diagonal structure shows some degree of category-level confusion, yet the dominant diagonal pattern confirms that stimulus-specific information is still reliably recovered. (B) Classifier transfer to the main task: classifier probabilities show a peak following stimulus presentation for shown items (blue) but not non-shown items (gray).

#### Replay results day 2 visual cortex before structural change

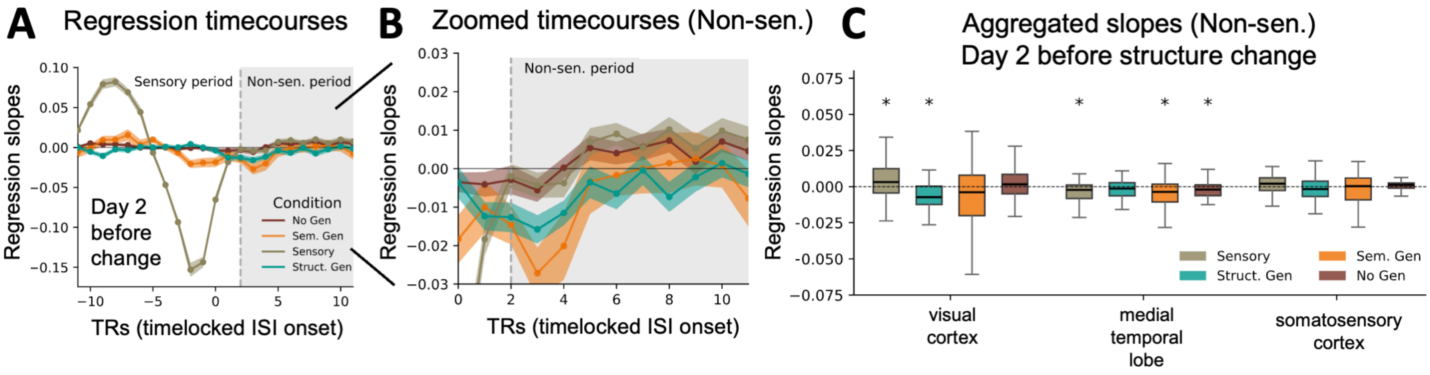


Figure 7: **Replay evidence in visual cortex on Day 2 before the structural change.** (**A–B**) Regression timecourses timelocked to ISI onset show replay dynamics during the sensory and non-sensory periods. The pattern closely resembles Day 1 (Fig. 5 in the main text): during the sensory period, activity tracks the currently observed stimulus, and during the subsequent non-sensory period, the reward-linked sequences are reactivated. (**C**) Aggregated regression slopes from the non-sensory period reproduce the main characteristics observed on Day 1. Statistical testing here is based only on the first two blocks of Day 2 (i.e., before the hidden structure change) and is therefore less powered, but the overall replay profile remains consistent with the main-text results.
